# Redox-active compound generated by bacterial crosstalk induces hypha branching in *Streptomyces* species

**DOI:** 10.1101/2023.01.12.523877

**Authors:** Manami Kato, Shumpei Asamizu, Hiroyasu Onaka

## Abstract

Chemical cross talks between *Mycolicibacterium septicum* HEK138M and *Bacillus subtilis* 168 affect the bacterial morphology of *Streptomyces variegatus* HEK138A. We found that *S. variegatus* exhibits unusual hyphae branching by the bacterial interaction. We aimed to elucidate the mechanism by performing activity guided purification of substances that induce the unusual cell morphology. We found that pyrogallol, a redox active aromatic small molecule induced significant hyphae branching in *S. variegatus* and the activity was also observed in some of other *Streptomyces* species. Interestingly, the pyrogallol activity was diminished by adding catalase, which broke down H_2_O_2_. To further confirm the involvement, H_2_O_2_ was tested and similar activity which induced hyphal branching was observed. This indicates that reactive oxygen species (ROS) generated by redox-active compound (RAC) is the inducing factor of hyphae branching. Further investigation revealed that pyrogallol was generated by NahG enzyme homolog of *M. septicum* using 2,3-dihydroxybenzoic acid as substrate by heterologous expression in *E. coli*. Moreover, co-culture with gene knock-out mutants revealed that 2,3-dihydroxybenzoic acid was supplied by *B. subtilis* produced as intermediate of bacterial siderophore bacillibactin. Since the hyphae branching of vegetative mycelium can increase the density of filamentous network and consequently help secure the milieu in soil, our results suggested that those filamentous soil bacteria use ROS which can be supplied from plant derived RAC as a signal. As those RAC ubiquitously exist in soil environment, the system will be beneficial for sensing the nutrient sources in addition to the generally considered defensive response to oxidative stress.

**Importance:** The characterization of interactions between three or more bacteria are lacking as these interactions are visually imperceptible in general. Our current study revealed changes of morphological behavior by the bacterial interaction. This study showed that hydrogen peroxide generated by redox-active compound derived from a breakdown product of siderophore can significantly increase the number of hyphae tip extension in filamentous bacteria. Our result implies the existence of oxidative response system using a low amount of reactive oxygen species as an integrated signal to sense the plant-derived carbon source by the filamentous soil bacteria. As a result of sensing, filamentous soil bacteria may decide whether the hypha tip should be extended to further explore the area or increase the tips to densify filamentous network to monopolize the nutrients in the milieu.

## Introduction

*Streptomyces* species is an industrially important Gram-positive filamentous actinobacteria, which produces bioactive natural products (secondary metabolites: SMs) that can be used as antibiotics, pesticides, and anti-cancer drugs (1, 2). Actinomycetes including *Streptomyces* species can interact with mycolic acid-containing bacteria (MACB) such as *Tsukamurella pulmonis* TP-B0596; such bacterial interactions can have effects on the production of specialized metabolites by actinomycetes, refer to as combined-culture (3–5). MACB can activate the production of undecylprodigiosin in *Streptomyces lividans* TK23, which are not produced by this strain under general laboratory culture conditions (3, 6). Although the mechanism of induced production of SMs is yet to be elucidated (7), interactions among *Streptomyces* species and MACB likely occur naturally since the combination of these two species has been co-isolated from environmental soil (8). These observations evoke interests around the function of bacterial (secondary) metabolism for constructing the microbial community in a soil environment where *Streptomyces* species and MACB exist simultaneously.

MACB are a group of actinomycetes that contain specific long-chain fatty acids (C_30_–C_60_ mycolic acids) in the cell wall (9). MACB has intrinsic resistance to antibiotics, which is conferred by the characteristic mycobacterial cell wall to avoid the penetration by antibiotics (10). Although the profit for *Streptomyces* species in this combination is obscure, these properties of MACB, inducer of SMs and resistance to antibiotics, can be considered to be responsible for the “combined” action of *Streptomyces* bacteria and MACB to fight against the third party. Previous studies have reported several cases of pairwise interactions using SMs that involve additional third party (11, 12). For example, the association of fungus-growing leafcutter ants with actinomycetes has been previously reported. Leafcutter ants cultivate fungal gardens (of *Leucoagaricus* sp.) as a food source, but they are constantly threatened by invasive fungal species, such as *Escovopsis* sp. To compete against the invasive fungi, the ants also carry actinomycetes associates that produce antifungal SMs (e.g., candicidin D, nystatin, and dentigerumycin). Knowledge of visually perceptible interactions among more than three organisms is relatively well accumulated, however the interactions among three or more bacteria are largely unexplored as they are generally imperceptible in a cell size scale.

*Streptomyces* species have a complex life cycle unlike other bacteria that undergo binary fission. Their life cycle begins with a spore, and its germination initiates with the emergence of a germ tube. The germ tube extends by filamentous cell, which extends using polar tip to yield long hyphal filaments, and occasionally hyphal branches emerge from the trunk filament. This cell growth manner with the branching of hyphae leads to the formation of a filamentous cell network in vegetative growth phase (13, 14). Tip extended filamentous cells further undergo cell differentiation to aerial mycelium and then proceed cell division forming spore to complete life cycle. Unique example has been known such as exploratory growth of *Streptomyces* species by significant polar tip extension, which is induced by alkalized condition caused by fungi (15). However, as *Streptomyces* species has evolved complex response system for diverse stimuli that can be derived from environment including bacterial interaction (16), what we have observed so far for morphological behavior would be only a limited fraction and large parts are remain unrevealed.

In a previous study, we found a natural combination of *Streptomyces* species and MACB that were co-isolated from environmental soil collected at Hegura Island, which involved alteration of secondary metabolism (8). In the combination that was retrieved, extracts from combined-culture among *S. variegatus* HEK138A and MACB: *Mycolicibacterium septicum* HEK138M showed specific antibacterial activity against Gram-positive bacteria, such as *Bacillus subtilis*. In the present study, we aimed to investigate the intergeneric interaction among *Streptomyces* species, MACB, and *B. subtilis*. By analyzing the phenotypic changes in the interaction area using scanning electron microscopy, we found abnormal cell shape formed in mycelia of *S. variegatus* HEK138A. Here, we report the characterization of metabolic interactions that generate redox active small molecule, which cause the induction of the disordered hyphal branch in *Streptomyces* species.

## Results

### Disordered hyphae branching of S. variegatus by bacterial interaction

*S. variegatus* HEK138A and *M. septicum* HEK138M was co-isolated from soil sample collected in Hegura island, Japan (8). Extracts from the combined-culture of these two species showed specific antibacterial activity against Gram-positive bacteria such as *B. subtilis* (8). We developed dual culture of *S. variegatus* and *M. septicum* to investigate whether the specific growth inhibition of *B. subtilis* can be observed on agar plate, as has been previously reported for the other combinations (5). In addition, we also investigated the phenotypic changes in the interaction area by scanning electron microscopy (SEM) to capture changes that were visually imperceptible. We noticed the slight color change at the edge of *S. variegatus* colony on agar plate culture (Fig. 1a). The cell morphological differentiation (aerial mycelium formation or spore formation) in *S. variegatus* was likely induced by chemical modulators that were produced as a result of bacterial interactions. This speculation was based on the fact that some natural products are known to change the cell differentiation of *Streptomyces* species (17–19). However surprisingly, the SEM image of the *S. variegatus* mycelium at the interaction area did not show the cells of developed morphology with aerial mycelium; instead, they showed irregular form of the vegetative mycelium (Fig. 1b). The mycelium observed using SEM showed disorderly emergence of hyphae branches; significant number of hyphae tip emerged from trunk mycelia (Fig. 1c-d).

**FIG 1.**
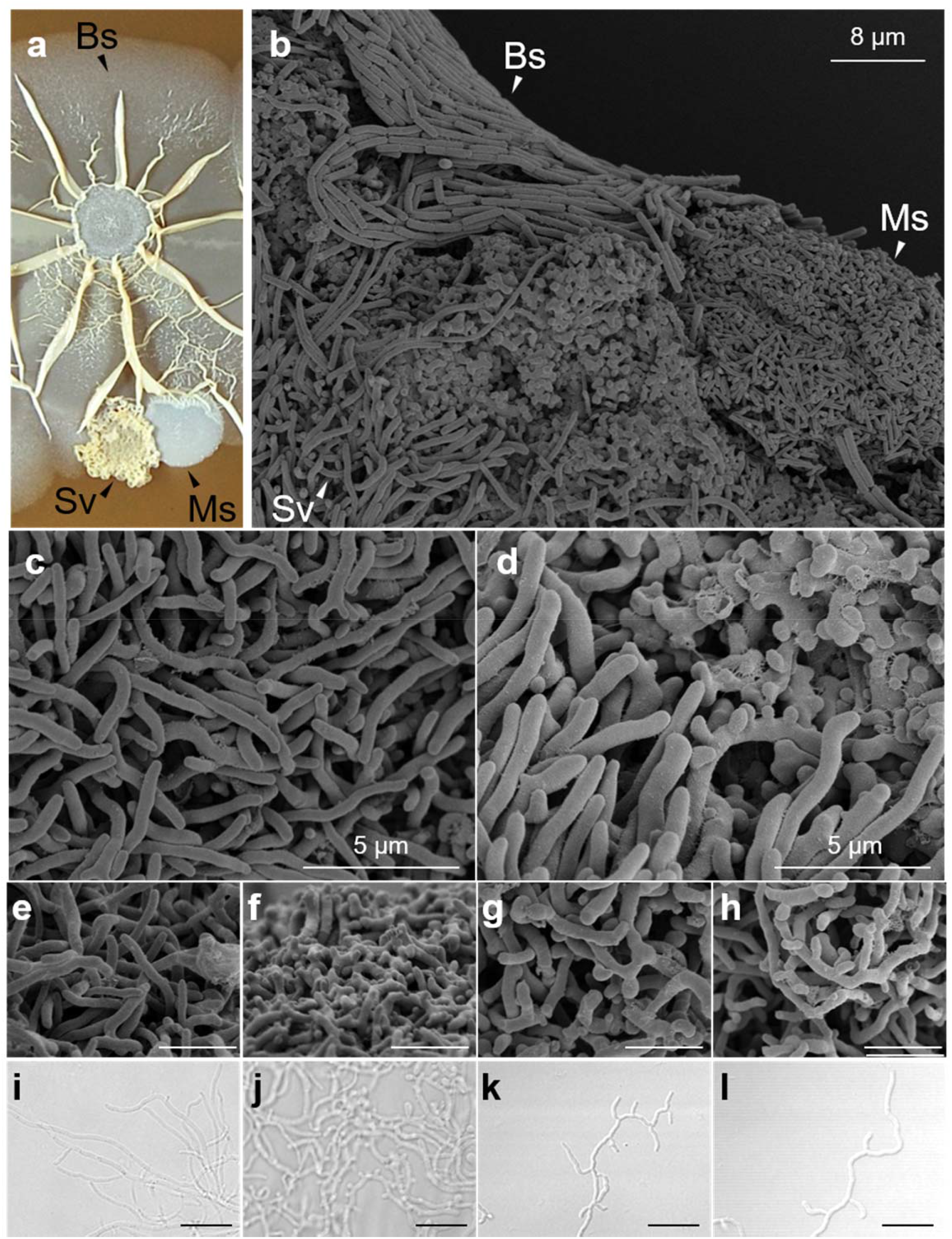
Disordered hyphae branching of *S. variegatus*. (a) Triple-culture of Sv, Ms, and Bs on agar plate. (b) SEM image of the interaction area of Sv, Ms, and Bs. (c) Magnified SEM image of normally grown Sv mycelium on agar plate. (d) Magnified SEM image of Sv mycelium at the interaction with Ms and Bs on agar plate. SEM images of colony (e-h) and DICM images of mycelium grown in slide culture (i-l). Morphology of normally grown *S. variegatus* cell (e, i); morphology of *S. variegatus* cell by addition of butanol extract from triple-culture (f, j); by addition of authentic pyrogallol (g, k); and by addition of H_2_O_2_ (h, j). Sv: *S. variegatus* HEK138A; Ms: *M. septicum* HEK138M; Bs: *B. subtilis* 168. (scale bar: 5 μm for a-d and 10 μm for e-h)

### Small molecules generated by bacterial interaction induce disordered hyphae branching

We first investigated which bacteria of two (*M. septicum* or *B. subtilis*) can induce the formation of disordered hyphae branching on mycelium of *S. variegatus*. Interestingly, the competitive dual culture did not induce the disordered hyphae branching on the mycelium of *S. variegatus* that was comparable to those of the triple-culture (Fig. S1). This indicates that the induction of disordered hyphae branching require the interactions between both *M. septicum* and *B. subtilis*. Thereafter, we tested whether organic substance(s) can induce the disordered hyphal branching of *S. variegatus* mycelium. For this, the triple-culture (solid) of *S. variegatus, M. septicum*, and *B. subtilis* was extracted using butanol and spotted next to the colony of *S. variegatus* that was grown on agar plate. After incubation, the *S. variegatus* colony was fixed for SEM imaging. The SEM image showed that substance(s) that was extracted using butanol can induce the emergence of similar disordered hyphae on the mycelium similar to those observed at the bacterial interaction area (Fig. 1f). We additionally analyzed the activity using slide culture, and the induction of hyphae branch by the culture extracts was also observed under a differential interference contrast microscope (DICM) (Fig. 1j). It was first presumed that bacitracin produced by *B. subtilis* was involved in the disordered hyphae emergence on mycelium of *S. variegatus* (20). However, this possibility was eliminated as *B. subtilis* 168 used in the experiment lacked bacitracin biosynthetic gene cluster and cannot produce bacitracin (21). Hence, in the present study, we performed purification to identify the active substance(s) that induce the hyphal branching.

### Bioactivity guided isolation of substance that induce disordered hyphae branching

Triple-culture (liquid) of the three strains were extracted using ethyl acetate and the active substances were further purified using column chromatography which was guided by hyphae branch inducing activity (Fig. S2 shows the purification scheme). The activity was confirmed by images of the colony obtained by SEM or those of slide culture under DICM. Interestingly, when the culture extracts were subjected to Sephadex LH20 column chromatography, activity was observed in two different fractions (No. 7 and 9) (Fig. S3). Both active fractions were further subjected for purification, but low yield and poor chemical signature of the substance(s) from the fraction No. 9 prevented further isolation of the compound. Therefore, we focused on the substance obtained from fraction No. 7 and further purified by open ODS column and preparative high-performance liquid chromatography (HPLC), and successfully isolated the active compound. The spectral data (HR-ESI-QTOF-MS, ^1^H NMR, and ^13^C NMR) of the purified compound contained chemical shifts that were identical to a compound known as pyrogallol. ^1^H NMR: 6.31 (2H, d, *J*=8.0), 6.49 (1H, t, *J*=8.2); ^13^C NMR: 147.3, 134.5, 120.2, 108.4; HR-MS: obs. 127.0395 [M+H]^+^ (calc. 127.0390 [M+H]^+^ for molecular formula of C_6_O_3_H_7_^+^) (Fig. S4, Table S1). Hence, commercially available pyrogallol was used for further experiments.

### Effect of pyrogallol against S. variegatus, M. septicum, and B. subtilis

Pyrogallol is known as inhibitor of bacterial growth (22). Hence, we first determined the concentration of pyrogallol that show the growth inhibition using agar medium assay. We found that 32 μg/ml of pyrogallol can completely inhibit the growth of *S. variegatus* and *M. septicum; B. subtilis* showed relatively high tolerance to pyrogallol (inhibition concentration: 256 μg/ml) (Table S2). Subsequently, different concentrations of pyrogallol (0.1, 1, and 10 μM) were tested against *S. variegatus* and activity of hyphae branching was observed under DICM. Obtained image confirmed that pyrogallol can consistently and reproducibly induce the hyphae branching (Fig. S5a). However, when the SEM images of the bacterial interaction area and of the area in presence of pyrogallol were compared (Fig. 1d and 1g), slightly different images were obtained. The mycelium at the interaction area was rich in saccular-like structure with smooth surface (Fig. 1d). In contrast, the presence of pyrogallol resulted in rough surface with rather deformed cells (Fig. 1g). This difference in morphology maybe attributed to the presence of at least two substances in the culture extract that can induce the abnormal cell form (Fig. S3). Thus, mixture of these substances may be responsible for the cell morphology observed at the interaction area. Although the SEM image showed slightly different morphology, slide culture images showed the clear and significant emergence of hyphae branch (Fig. 1k and S4a), strongly indicating that addition of pyrogallol induce hyphae branching.

### Pyrogallol induced hyphae branching in a broad range of Streptomyces species

Subsequently, to investigate whether pyrogallol can induce hyphae branching in other *Streptomyces* species, same concentrations of pyrogallol was tested for six other *Streptomyces* species that are generally used for biological experiments. Similar to the mycelium of *S. variegatus*, the mycelia of *S. griseus* IFO13350, *S. albus* J1046, and *S. lividans* TK23 showed significant emergence of hyphae branch in all concentrations consistently and reproducibly (Fig. S5b-d). *S. venezuelae* NBRC13096 and *S. avermitilis* MA4680 receive similar effect, but was weaker than that observed in *S. variegatus* (Fig. S5e-f). The tested concentration did not show the expected phenotypic changes in *S. coelicolor* A3(2) M145 (Fig. S5g); however, the reason is not clear yet. Additionally, effects of pyrogallol (10 μM) for *M. septicum* and *B. subtilis* was also tested, which showed that most of the *B. subtilis* cells showed ordinary shape, but several cells also showed round shape rather than rod shape (23) (Fig. S6a), and *M. septicum* sustained no apparent phenotypic change (Fig. S6b).

### Quantification of lengths of tip-to-branch and branch-to-branch

As *S. griseus* showed similar response by addition of pyrogallol to *S. variegatus* (Fig. S5a-b), We further used *S. griseus* to evaluate the quantitative significance of the inducing activity of hyphae branch by pyrogallol. AcGFP1 was expressed in *S. griseus (S. griseus::Acgfp1*) to illuminate the mycelium. The images of slide culture were obtained under confocal laser scanning microscope (CLSM). We used fluorescence excitation (488 nm) and emission (510 nm) wavelength to confirm that AcGFP1 is expressed in every mycelium. It was observed that mycelium cells were thoroughly illuminated by fluorescence AcGFP1 (Fig. S7). After obtaining the multiple images of CLSM, we performed image extraction, disassemble process, and length measurement using ImageJ (Fiji), which was employed in JAVA-based WEKA platform (24). We disassembled the image of the grown hyphae, and lengths of the tip-to-branch and branch-to-branch were measured to obtain the quantitative data. Lengths of the tip-to-branch and branch-to-branch were used for criterion to define the frequency of hyphae branch. The addition of 1 μM pyrogallol significantly reduced the length of tip-to-branch (0.71-fold) by comparing the median value (3.08 nm by addition of pyrogallol and 5.32 nm in control) (p=4.34×10^-9^) (Fig. 2b-c and 2g left). Additionally, the length of branch-to-branch also reduced significantly (0.71-fold) by comparing the median value (8.17 nm by addition of pyrogallol and 11.47 nm in control) (p=1.27×10^-7^) as well (Fig. 2g right). The result confirmed that pyrogallol induces the generation of short mycelium in high frequency.

**FIG 2.**
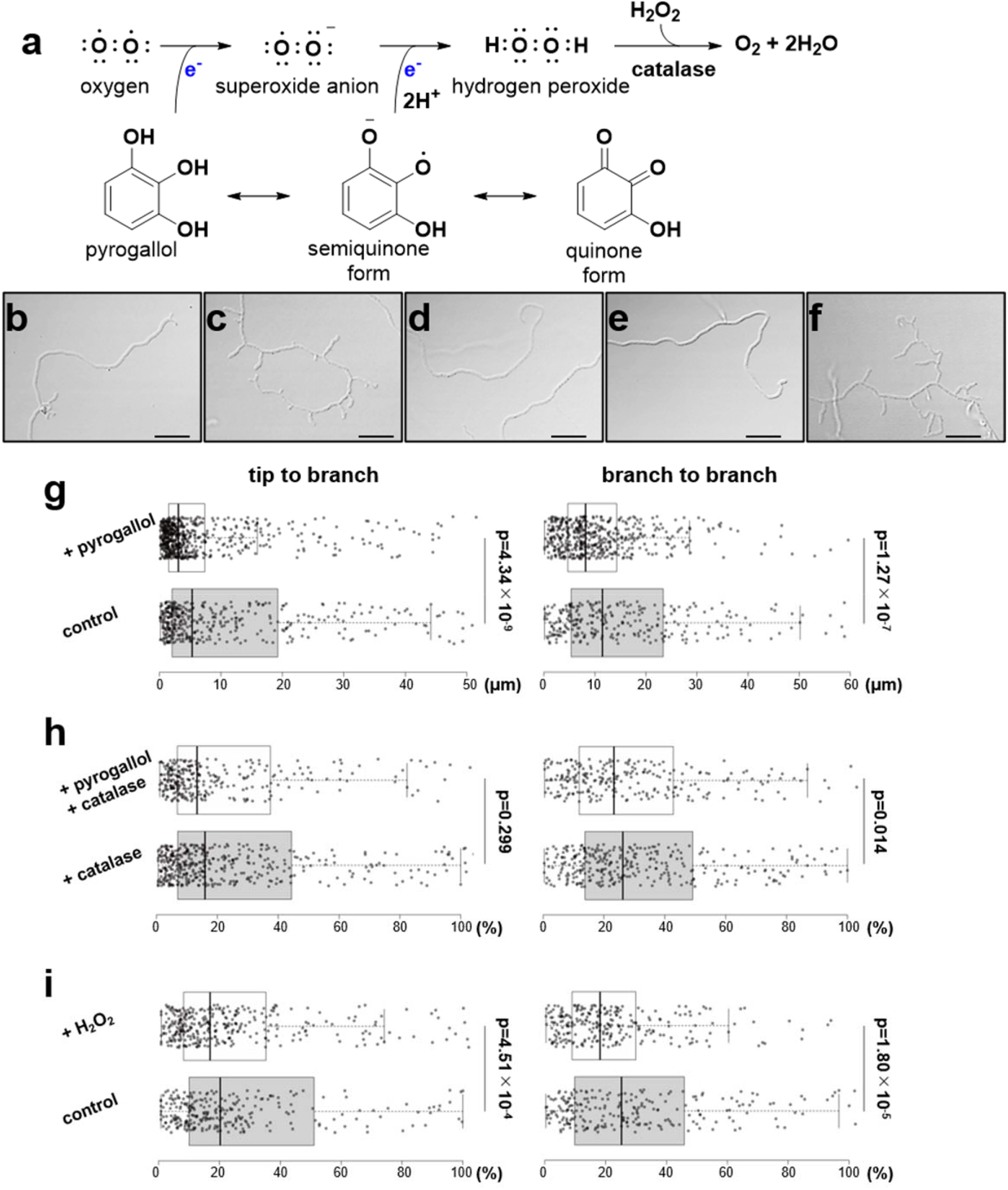
Disordered hyphae branching of *S. variegatus* induced by H_2_O_2_ generated from RAC. (a) Reaction scheme of pyrogallol (RAC) to generate H_2_O_2_ (ROS). (b-e) DICM images of *S. griseus* IFO13350 (AcGFP1) mycelium grown in slide culture. Normally grown cells (b); Cell morphology by addition of 1 μM pyrogallol (c); by addition of 1 μM pyrogallol and 22 U/ml catalase (d); by addition of 22 U/ml catalase (e); by addition of 0.88 μM H_2_O_2_ (f). (scale bar = 10 μM) (g-i) Box-and-whisker plot of tip-to-branch and branch-to-branch lengths. Effects by pyrogallol (1 μM) (g); Effects of pyrogallol (1 μM) with catalase (22 U/ml) (h); Effects of H_2_O_2_ (0.88 μM) (i). Center lines show the medians; box limits indicate the lower and upper quartile; whiskers extend to 5 and 95 %. Data points are plotted as dots (n=422, 616 sample points).

### Hydrogen peroxide generated by pyrogallol induced hyphae branching

Catechol (*ortho*-hydroxyphenol) or quinol (*para*-hydroxyphenol) is well characterized as RAC, which can react with molecular oxygen to be oxidized and form (*ortho*- or *para*-) quinone, and readily generate oxide such as hydrogen peroxide (H_2_O_2_) (25). Therefore, we hypothesized that the H_2_O_2_ can induce the emergence of hyphae branching in mycelium of *Streptomyces* species since chemically similar effect can be applied for pyrogallol (Fig. 2a) (26). To test this hypothesis, we first added catalase to the culture medium containing 1 μM pyrogallol to breakdown the generated H_2_O_2_. Catalase can convert two molecules of H_2_O_2_ to two molecules of H_2_O and one molecule of O_2_ to detoxify the oxidative stress (Fig. 2a) (27). *S. griseus::Acgfp1* was cultured in catalase-containing medium with or without pyrogallol, and further hyphae lengths were measured. The result showed that the emergence of hyphae branch which was expected to be induced by pyrogallol was almost abolished by addition of 22 U/ml of catalase (Fig. 2d-e). The quantification analysis showed that lengths of both tip-to-branch and branch-to-branch have no significant differences in lengths in presence of catalase (p-value=0.299 and p-value=0.014, respectively) (Fig. 2h). The result revealed that activity of pyrogallol was quenched by catalase which decompose H_2_O_2_. To further verify the involvement of H_2_O_2_ as an inducer of hyphae branch, we then added H_2_O_2_ to the culture of *S. griseus::Acgfp1*. By adding 0.88 μM of H_2_O_2_, we notably observed the similar phenotype comparable to addition of pyrogallol, which the emergence of hyphae was induced (Fig. 2b-c and 2f). The result of quantification supported the significant emergence of hyphae by the values that both length of tip-to-branch and branch-to-branch become shorter (0.84-fold and 0.69-fold, respectively) with significance (p-value=4.51 × 10^-4^ and p-value=1.80 × 10^-5^, respectively) by addition of H_2_O_2_ (Fig. 2i). The result revealed that H_2_O_2_ generated by pyrogallol is the signal to induce emergence of hyphae branch.

### Pyrogallol was generated by M. septicum from 2,3-DHBA supplied by B. subtilis

To identify the producer(s) of the pyrogallol, we cultured *S. variegatus, M. septicum*, and *B. subtilis* in possible combinations and analyzed the extracted metabolites using HPLC equipped by photodiode array (PDA). PDA-HPLC traces showed that pyrogallol was produced in co-culture of *M. septicum* + *B. subtilis* (930 ± 40 μM) and triple-culture of *S. variegatus* + *M. septicum* + *B. subtilis* (100 ± 40 μM), indicating that *S. variegatus* is not directly involved in the production of pyrogallol (Fig. 3b and S8). Previous studies showed that gallic acid (28) or 2,3-DHBA (29) can be a direct precursor of pyrogallol by microbial conversion. Therefore, we tested the conversion by feeding gallic acid (0.5 mM) or 2,3-DHBA (0.25 mM) to the culture of *M. septicum* or *B. subtilis*. HPLC traces revealed that pyrogallol was formed only when 2,3-DHBA was fed to the culture of *M. septicum* with yield of 37.3 ± 9.8 μM (14.9% conversion rate) (Fig. 3c and S9). It is known that *B. subtilis* produces 2,3-DHBA as an intermediate of its siderophore, bacillibactin (30) (Fig. 3a). Hence, we hypothesized that 2,3-DHBA secreted by *B. subtilis* was supplied to *M. septicum* as a precursor of pyrogallol. To test this, we co-cultured *M. septicum* with knock-out mutants of the bacillibactin biosynthetic genes (*dhbA-F*) of *B. subtilis* (30, 31). When *M. septicum* was co-cultured with *B. subtilis ΔdhbF* or *ΔdhbE*, both of which can accumulate 2,3-DHBA as intermediate, pyrogallol was detected in the co-culture broth (Fig. 3d and S10). In contrast, when *M. septicum* was co-cultured with Δ*dhbC*, *ΔdhbB* or *ΔdhbA* strains which cannot accumulate 2,3-DHBA, pyrogallol was not detected (Fig. 3d and S10). Generation of pyrogallol was consistent with the availability of 2,3-DHBA in the blocked mutants. Notably, the result demonstrated that 2,3-DHBA, bacillibactin biosynthesis intermediate, was direct precursor of pyrogallol synthesis by *M. septicum*.

**FIG 3.**
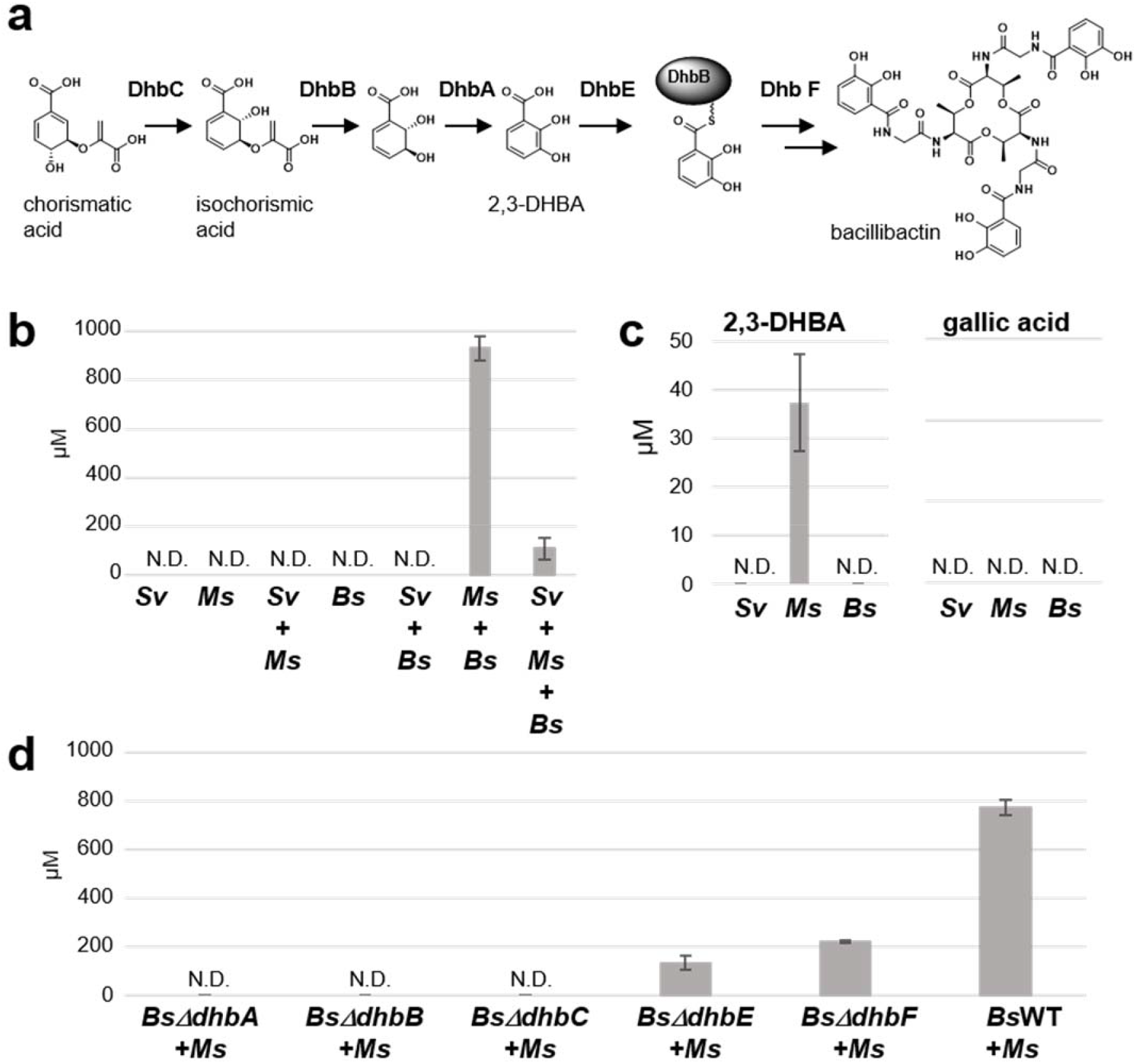
Biosynthetic origin of pyrogallol. (a) Biosynthetic pathway of bacillibactin in *B. subtilis*. (b) Production of pyrogallol in single-, dual- and triple-culture. (c) Feeding of 2,3-DHBA or gallic acid as potential precursor in the culture of Sv, Ms, and Bs. (d) Co-culture of Ms with bacillibactin biosynthetic gene disruptants (*ΔdhbA-F*) of Bs. Sv: *S. variegatus* HEK138A; Ms: *M. septicum* HEK138M; Bs: *B. subtilis* 168. (N.D.: not detected)

### Salicylate 1-monooxygenase (NahG) from M. septicum convert 2,3-DHBA to pyrogallol

As pyrogallol was synthesized from 2,3-DHBA by *M. septicum*, we searched the enzyme that convert 2,3-DHBA to pyrogallol. Previously, *nahG* (salicylate 1-monooxygenase, EC 1.14.13.1) from *Pseudomonas putida* DOT-T1E was reported to convert 2,3-DHBA to pyrogallol using *E. coli* as host (29). Draft genome sequence of *M. septicum* was obtained, and a possible NahG^Ms^ enzyme with 31% similarity in deduced amino acid were found (Fig. S11). The *nahG*^Ms^ was cloned and expressed under T7 promoter in *E. coli* BL21(DE3) (Fig. 4a). Using *E. coli* resting cell expressing the NahG^Ms^, we incubated 2,3-DHBA or salicylate (as native precursor) and traced the conversion using PDA-HPLC. As expected, generation of pyrogallol from 2,3-DHBA (Fig. 4b-e), as well as catechol from salicylate (Fig. S12) was observed only in the reaction with *E. coli* resting cell expressing the NahG^Ms^. Hence, it was revealed that NahG enzyme from *M. septicum* can accept 2,3-DHBA as well as salicylate as substrate and catalyze oxidative decarboxylation to produce pyrogallol or catechol.

**FIG 4.**
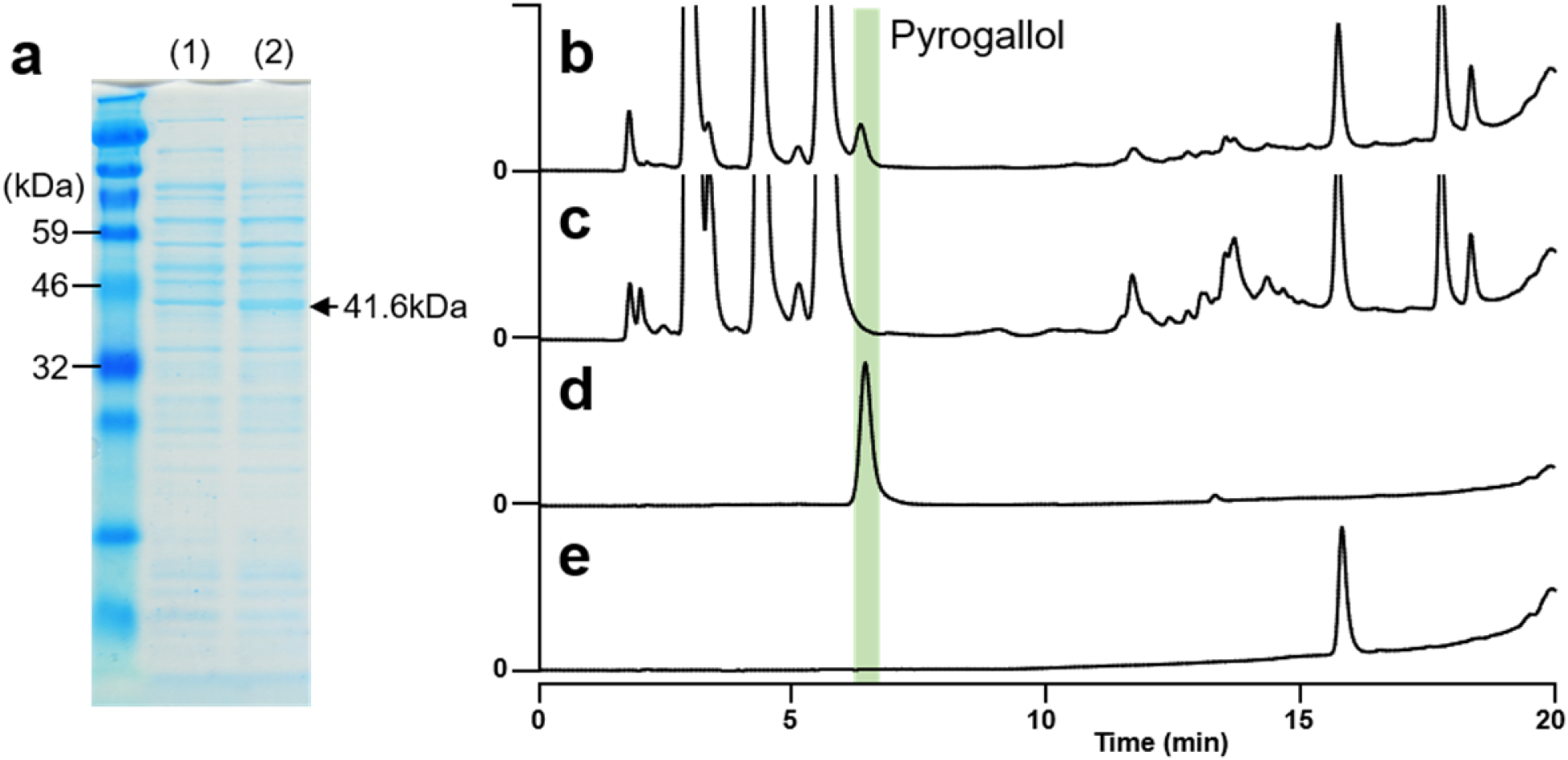
Bioconversion of 2,3-DHBA to pyrogallol by NahG^Ms^. (a) SDS-PAGE of soluble protein fraction form *E. coli* expressing NahG^Ms^ (2) and negative control with empty vector (1). (b-e) PDA-HPLC traces of bioconversion from 2,3-DHBA to pyrogallol. (b) Incubation of 2,3-DHBA with *E. coli* resting cell expressing NahG^Ms^. (c) Incubation of 2,3-DHBA with *E. coli* resting cell. (empty vector control) (d) Pyrogallol standard. (e) 2,3-DHBA standard.

### Estimation of the distribution of 2,3-DHBA and nahG homolog in soil environment

Several bacterial species are known to produce siderophore with catechol units, whose precursor is 2,3-DHBA (32). We searched for the orthologous gene set that is responsible for the production of 2,3-DHBA, which comprise *dhbC* (isochorismate synthase), *dhbB* (isochorismate lyase domain), and *dhbA* (2,3-dihydro-2,3-dihydroxybenzoate dehydrogenase) using KEGG database. We found that 10.6% (789/7393) of bacterial strains possessed the gene set, and remarkably, percentile was 84.7% (111/131) in *Bacillus* species. This bioinformatic result implies that 2,3-DHBA abundantly exist in the soil bacterial community. We then searched for the *nahG* ortholog gene in MACB (such includes *Tsukamurella, Rhodococcus, Corynebacterium, Gordonia*, and *Nocardia* spp.). It was found that 18.1% (28/155) of MACB possessed the *nahG* ortholog, and percentile was 10.5% (4/38) for *Mycolicibacterium* species. Overall, the result implied that our observation of significant emergence of hypha branch on mycelium of *Streptomyces* species by pyrogallol may stochastically occur in the environmental bacterial soil community.

## Discussion

Our study revealed that pyrogallol was a decomposed product derived from the assembly component for catechol-type siderophore. This metabolic interaction between MACB and *Bacillus* species likely represents the competition for acquiring iron from the environment. Our present study also identified that redox active compound (RAC) such as pyrogallol, can modulate cell morphology of *Streptomyces* species *via* generation of reactive oxygen species (ROS) such as H_2_O_2_ (Fig. 5). Interaction involving bacteria observed in our current study may stochastically occur in the soil environment, however, our observation also suggested the general behavior of filamentous bacteria against oxidative signals.

**FIG 5.**
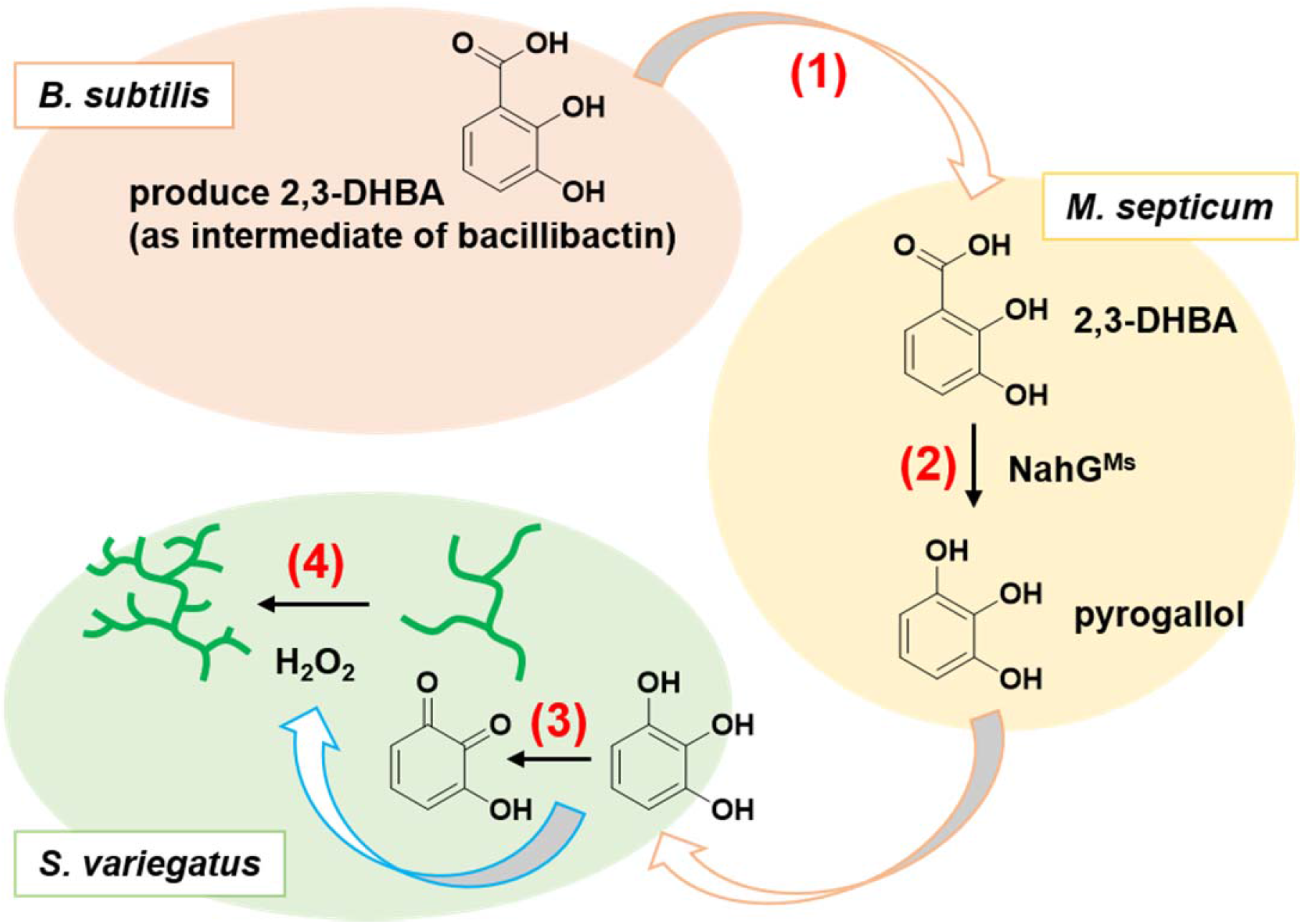
Summary of triple bacterial interaction. (1) *B. subtilis* secrete 2,3-DHBA as intermediate of siderophore bacillibactin. (2) *M. septicum* HEK138M convert 2,3-DHBA to pyrogallol by NahG^Ms^ (salicylate 1-monooxygenase). (3) pyrogallol convert to quinone-form and result in generation of H_2_O_2_. (4) H_2_O_2_ induce emergence of hyphae branch in *Streptomyces* species.

### SMs that affect the morphological behavior of Streptomyces species

Several SMs from *Bacillus* species have been reported to show intergeneric chemical communications between *Streptomyces* and *Bacillus* species. *B. subtilis* produces surfactin, which is a lipopeptide antibiotic that inhibits the formation of aerial hyphae of streptomycetes by defecting the hydrophobic cell wall (33). *B. subtilis* also produces bacillaene, NRP/PK hybrid compound, that inhibits the production of undecylprodigiosin by *S. coelicolor* A3(2) (34). As described before, bacitracin modulates the emergence of hyphae by inducing the phosphorylation of DivIVA protein (20). Additionally, several other examples of interspecies or intergeneric chemical cross talk have been previously reported. Desferrioxamines, which is bacterial siderophore described as public goods for ion acquisition, can be donated by a bacteria species; it affects the cell differentiation and antibiotic production of receiver *Streptomyces* species (35). Furthermore, several specific SMs have been reported to induce aerial mycelium, such as hormaomycin (19) (or takaokamyicin) (36) from *S. griseoflavus* W-384 and panamycin-607 (37) from *S. alboniger* IFO 12738. In sublethal concentrations, goadsporin (17) and lincomycin (38) affect both secondary metabolism and morphological differentiation in other actinomycetes. In addition to the previously known bacterial strain specific specialized SMs, our findings identified that pyrogallol which is more generally exist in the environment modulate cell morphology of *Streptomyces* species. This likely suggest that *Streptomyces* species has evolved unsolved response systems for more general stimuli in the environment.

### Signals for hypha branch in Streptomyces species

Hyphae branching of vegetative mycelium can be beneficial as it increases the density of filamentous cell network, which can help secure the milieu in soil environment. However, knowledge of the mechanisms how to determine the points of hyphal tip emergence (or polarisome) in *Streptomyces* species is still unknown. Hypha extension and hyphal branching involves two coiled-coil proteins, namely DivIVA (39) and Scy (40), which form protein complexes described as polarisomes as well as intermediate filament-like protein FilP (41) that interacts with the polarisome. The DivIVA is essential for recruiting the two coiled-coil proteins, whereas Scy colocalizes with DivIVA to construct scaffold in the polarisome and FilP provides mechanical support by forming a stress bearing network behind the hyphal tip. Partial or null deletion or overexpression of the genes can affect the tip extension and hyphae branching (42), indicating that modulation of the polarisomes can disturb the extension of polar tip of the hyphae. Excessive phosphorylation of DivIVA by a bacterial serine/threonine kinase AfsK can modulate the polar extension and branching, particularly in response to inhibitors of cell wall synthesis, such as bacitracin and vancomycin (20). Co^2+^-binding bacitracin binds tightly to the undecaprenyl pyrophosphate, which interferes with cell wall synthesis by preventing its hydrolysis to monophosphate, which is essential step in peptidoglycan biosynthesis (43). In addition, osmotic challenge can also induce hyphal branching in *S. coelicolor* M145 (44). However, mechanism which direct the emergence of hyphae in the high osmotic shift challenge is yet unknown. Although future studies should be required to identify the mechanism by which disturbance of polarisome is involved in disordered hyphae branching by H_2_O_2_, our present study newly demonstrated that H_2_O_2_ generated by RAC become a diving force for the emergence of hypha branch.

### Response system for ROS in Streptomyces species

Pyrogallol can generate ROS including H_2_O_2_ by conversion to its *ortho*-(benzo)quinone form. Recently, several SMs including actinorhodin (45) or granaticin (46) have been revealed as RAC that generate H_2_O_2_. Although the molecular response regulation of *Streptomyces* species in response to oxidative stress is relatively well characterized (47), ROS-induced emergence of hypha has not been described. The ability to respond to oxidative stress is critical for survival, and organisms have evolved diverse strategies to sense oxidative stress. When face with direct oxidative challenge in the environment, bacteria exercise a system to detoxify oxidants such as super oxide anion (O^2-^) and hydrogen peroxide (H_2_O_2_) by using two major regulatory protein, SoxR to induce superoxide dismutase (48) and OxyR to induce catalase (49), which is also conserved in *Streptomyces* species. The principal oxidative stress response in *Streptomyces* is controlled by a regulatory system consisting of a σ-factor SigR (47, 50), and a redox-sensitive anti-σ-factor, RsrA (51). The RsrA is a Zn^2+^ binding protein that binds to SigR with nanomolar affinity under reducing conditions, which prevents it from associating with RNA polymerase to activate transcription. When the cell encounters oxidative stress, an intramolecular disulfide bond forms between two of the cysteines, leading to expulsion of Zn^2+^ and inactivation of RsrA. Thereafter, SigR is released, which activates the transcription of a regulon consisting of more than 100 target genes. Regulon of SigR was investigated using ChiP-seq and revealed 108 regulons (52). In our search, the SigR regulon did not include genes involved in the hyphae tip extension such as DivVIA (*sco2077*), Scy (*sco5397*), FilP (*sco5396*), noncoiled-coil protein MreB (*sco2611*), FtsZ (*sco2082*), other coiled-coil protein *sco3114* and *sco2168*. Therefore, known principle of oxidative stress response involving SigR/RsrA or OxyR/SoxR in *Streptomyces* species may not directly be involved in the induced emergence of hyphae branch.

### Bacterial behavior in soil ecosystem

Though ROS are injurious by-products, they are essential participants in cell signaling and regulation in low concentrations (53). In soil, bacteria decompose plant components such as gallic acid, ellagic acid, tannic acid, and lignin that contain redox active groups (54), and those are used as carbon source for the growth of actinobacteria. Pyrogallol or catechol component which can generate low amount of H_2_O_2_ are decomposed products of those. Meanwhile emergence of hyphae branching can be considered to increase the density of filamentous network of *Streptomyces* species. Using low amount of H_2_O_2_ as an integrated signal to sense (recognize) the plant derived carbon source can be beneficial for the filamentous bacteria. Existence of those system can likely help decide whether to extend tip to explore more area or to increase the density of filamentous network to construct exclusive area which can take advantage of *Streptomyces* species to monopolize the nutrients.

## Materials and methods

### General

*S. variegatus* HEK138A and *M. septicum* HEK138M were isolated from Hegura Island in Japan, and the taxonomy of the strains were identified based on the 16S rDNA sequences (8). The culture conditions of *S. lividans* TK23, *S. griseus* IFO 13350, *S. avermitilis* MA-4680, *S. coelicolor* A3(2) M145 *S. venezuelae* NBRC13096, and *S. albus* J1046 followed the standard protocol (55). *B. subtillis* 168 and the BKE library (BKE32000, BKE31970, BKE31990, BKE31980, BKE31960) used in the present study were obtained from National BioResource Project (NIG, Japan). Additional information can be found in *Supplemental Material*.

## Acknowledgments

This research was supported by a grant-in-aid from the Institute for Fermentation, Osaka, (IFO) and Amano Enzyme Foundation (to S.A. and H.O.). This study was also partially supported by KAKENHI research grants (18K05385 to S.A. and 18H02120 to H.O.), general research grant from IFO (to S.A.), general research grant from Noda institute for scientific research (to S.A.), and the JSPS A3 Foresight Program. M.K. is supported by JST SPRING GX program. We thank Prof. Yasuo Ohnishi and Dr. Takeaki Tezuka for the AcGFP1 plasmid and technical supports in *S. griseus* genetic experiments. *Bacillus subtillis* BKE library were obtained from National BioResource Project (NIG, Japan). All authors declare no conflicts of interest associated with this manuscript. All authors designed research and analyzed data. M.K. performed all experiments. S.A. wrote the main text, M.K. and S.A. prepared figures and tables, and H.O. review the manuscript.

