## Supplemental material file for "Redox-active compound generated by bacterial crosstalk induces hypha branching in *Streptomyces* species"

Running Head: RAC induced hypha branching in *Streptomyces* spp.

**Abstract**

Chemical cross talks between *Mycolicibacterium septicum* and *Bacillus subtilis* 168 affect the bacterial morphology of *Streptomyces variegatus* HEK138A. We found that *S. variegatus* exhibits unusual hyphae branching by the bacterial interaction. We aimed to elucidate the mechanism by performing activity guided purification of substances that induce the unusual cell morphology. We found that pyrogallol, a redox active aromatic small molecule induced significant hyphae branching in *S. variegatus* and the activity was also observed in some of other *Streptomyces* species. Interestingly, the pyrogallol activity was diminished by adding catalase, which broke down $H_2O_2$. To further confirm the involvement, $H_2O_2$ was tested and similar activity which induced hyphal branching was observed. This indicates that reactive oxygen species (ROS) generated by redox-active compound (RAC) is the inducing factor of hyphae branching. Further investigation revealed that pyrogallol was generated by NahG enzyme homolog of *M. septicum* using 2,3-dihydroxybenzoic acid as substrate by heterologous expression in *E. coli*. Moreover, co-culture with gene knock-out mutants revealed that 2,3-dihydroxybenzoic acid was supplied by *B. subtilis* produced as intermediate of bacterial siderophore bacillibactin. Since the hyphae branching of vegetative mycelium can increase the density of filamentous network and consequently help secure the milieu in soil, our results suggested that those filamentous soil bacteria use ROS which can be supplied from plant derived RAC as a signal. As those RAC ubiquitously exist in soil environment, the system will take advantage for sensing the nutrient sources in addition to the generally considered defensive response to oxidative stress.

**Table of contents**

**Supplemental Methods**

***Culture conditions***

***Microscopy***

***Production, purification and structure elucidation of pyrogallol***

***HPLC analysis (for quantification of pyrogallol)***

***Construction of strain expressing AcGFP1 in S. griseus***

***Image extraction of hyphae branching and processing by Fiji (ImageJ)***

***Precursor (gallic acid or 2,3-DHBA) feeding experiment***

***Bioconversion of 2,3-DHBA to pyrogallol***

***Heterologous expression of nahG from M. septicum***

**FIG S1** Scanning electron microscope (SEM) image of cell morphology in competitive dual culture.

**FIG S2** Purification scheme of active compound.

**FIG S3** SEM images of *S. variegatus* HEK138A mycelia treated by active fractions.

**FIG S4** NMR and MS analysis of the isolated active compound.

**TABLE S1** NMR assignments.

**TABLE S2** MIC of pyrogallol against tested microorganism.

**FIG S5** DICM images of *Streptomyces* spp. mycelium treated with 0.1-10 μM of pyrogallol.

**FIG S6** DICM images of *B. subtilis* cells (a) and *M. septicum* HEK138M (b) treated with pyrogallol.

**FIG S7** DICM and CLSM images of *S. griseus* expressing AcGFP1 treated with pyrogallol.

**FIG S8** PDA-HPLC traces of metabolites in combination of all mixed-culture.

**FIG S9** PDA-HPLC traces of 2,3-DHBA (a) or gallic acid (b) feeding experiment.

**FIG S10** PDA-HPLC traces of co-culture between Ms and *dhb* gene knockout mutants.

**FIG S11** ClustalW alignment of NahG from *Pseudomonas putida* and *M. septicum* HEK138M.

**FIG S12** Bioconversion of salicylic acid to catechol by *E. coli* resting cell expressing $NahG^{Ms}$.

**TABLE S3** Oligo-DNA primers used for PCR in this study.

**References**

***Supplemental methods***

***Culture conditions***

**Agar plate competitive assay:** A3M agar medium (5 g glucose, 20 g glycerol, 20 g starch, 15 g pharma media, 3 g yeast extract, 20 g agar, and pH 7.0 in 1 L $H_2O$) was used. For competitive dual culture, 2 µL each of *S. variegatus* HEK138A spore stock ($1\times10^9$ CFU in 20% glycerol) and *M. septicum* HEK138M cell stock ($1\times10^{10}$ CFU in 20% glycerol) were inoculated by spots on the agar medium with space of 2 mm between the spots. For triple culture of *S. variegatus*, *M. septicum*, and *B. subtilis*, we further inoculated 2 µL of *B. subtilis* cell stock ($1\times10^7$ CFU in 20% glycerol), which were placed at a distance of 2 cm away from the spots on the agar medium. The cells were grown by incubation at 30°C for 5 days to observe the phenotype by scanning electron microscopy (SEM). **Slide culture:** We used 1 mm thickness of YGGS agarose medium (5 g glucose, 20 g glycerol, 20 g starch, and 3 g yeast extract, 20 g low melting point agarose (gelling temperature 30–31 °C), and pH 7.0 in 1 L $H_2O$) was prepared using glass plate (for SDS-PAGE gel) and was further used for slide culture. Before inoculation, spores of *Streptomyces* species were germinated by incubating the spore stock ($1\sim3\times10^4$ CFU) in 1 ml of ISP2 medium (Yeast extract 4 g, Malt extract 10 g, Glucose 4 g, in 1 L $H_2O$) at 30°C for 2 hrs. The germinated spores were used to inoculate on the edge of the slide culture. **Growth inhibition assay by pyrogallol:** To prepare the YGGS agarose medium, pyrogallol was first dissolved in $H_2O$ (3.2 mg/ml) and filter sterilized (0.2 µm filter). We added 0.6 ml of pyrogallol solution (2-fold dilution from 0.5 ng/ml to 320 mg/ml) to 6 ml of YGGS agarose. **Catalase assay:** Prior to casting slide culture agarose gel, 50 µl of 12.6 µg/ml (100 µM) pyrogallol and 100 µL of 100 µg/ml catalase (110 Unit) was added into 5 ml of YGGS agarose. For slide culture, 2 µl spore stock of *S. griseus* AcGFP1 (described later) (500~1000 cfu) was inoculated and further incubated at 30°C for 5 days. Hyphae branching was compared in the presence or absence of catalase solution. **$H_2O_2$ assay:** Prior to casting slide culture agar gel 30% $H_2O_2$ solution was diluted by distilled water to $3\times10^{-4}$% and 50 µL was added into 5 ml of YGGS agarose (880 nM $H_2O_2$ in final concentration). For slide culture, 2 µl spore stock of *S. griseus* AcGFP1 (described later) (500~1000 cfu) was inoculated and further incubated at 30°C for 5 days. Hyphae branching in the presence of $H_2O_2$ solution was compared with that in the absence of $H_2O_2$ solution.

***Microscopy***

To obtain SEM images, samples were prepared, as described here. Agar piece was prepared by cork borer and the sliced piece was transferred into 1.5 ml tube, and 1 ml of 2.5% glutaraldehyde in 0.1 M sodium phosphate buffer (pH7.4) was added and incubated for 1 hr at 4°C. After discarding the solution, the sample was rinsed using 1 ml of 0.1 M sodium phosphate buffer (pH7.4). Then, 1% $OsO_4$ in 100 mM sodium phosphate buffer (pH7.4) was added and incubated for 1 hr on ice. After discarding the solution, 1 ml of 0.1 M sodium phosphate buffer (pH7.4) was added to rinse the sample. Subsequently, the sample was dehydrated by stepwise addition of 50%, 70%, 90%, and 100% ethanol (1 ml each), and finally substituted by t-butyl alcohol, and stored at 4℃ until further use. Sample was dried under vacuum before Pt/Pd sputter coating (E-1030, Hitachi, Japan) and observed using scanning electron microscope (S-4800, Hitachi, Japan). The differential interference contrast microscope (DICM) images of the slide cultures of *Streptomyces* species were obtained by Olympus SZX16 equipped using a UPlanFLN 40×0.75 NA dry lens or UPlanFLN 100×1.30 NA oil-immersion lens (Olympus) and DP-20 digital camera (Olympus). Stereomicroscope images of competitive culture on agar plate were obtained using Nikon SMZ25 that was equipped with a SHR Plan Apo 1.6× lens (Nikon) and DS-Ri2 digital camera (Nikon). The confocal laser scanning microscope (CLSM) images of the slide culture of AcGFP1 in *S. griseus* were obtained using Olympus FV1200 equipped with a UPlanApo 100×1.40 NA oil-immersion lens (Olympus), they were observed at wavelengths of 488 nm (excitation) and 510 nm (emission) for AcGFP1. The Fluoview or FV10- ASW software was used for constructing images.

***Production, purification and structure elucidation of pyrogallol***

For the isolation of active compounds, *S. variegatus* and *M. septicum* was precultured using ISP2 medium (10 ml in test tube) at 30℃, 180 rpm for 2 days. *B. subtillis* was precultured by Miller Hinton medium (10 ml in test tube) at 30℃, 180 rpm for 2 days. Thereafter, 1 mL of each preculture were transferred into 100 ml of A3M medium in K-1 flask and further cultured at 30℃ at 200 rpm for 5 days. Equal volume of ethyl acetate (2.4 L) was added to the culture broth and extracted for 2 h. After centrifugation at 12,000 rpm for 15 min, ethyl acetate layer was collected and evaporated to obtain 3.2 g of crude extracts. This extract (3.2 g) was dissolved in 10 ml of DMSO and applied on Sephadex LH-20 open column (Sigma, 40 id × 1000 mm) in methanol. The extracts were eluted using methanol at a flow rate of 8 ml/min for total 1.8 L, and thereafter, the fractions were collected after every 200 ml. For bioassay, 5 ml were collected in part from each fraction and evaporated separately. The crude from 5 ml was dissolved in 200 µL of DMSO and 2 µL was used for bioassay using SEM. Two fractions (No. 7 and No. 9) contained active compound(s) and proceeded for further purification. The active crude extracts from fraction No. 7 were dissolved in DMSO and applied on Cosmosil C18-OPN column (Nacalai, 40 id × 300 mm). The extracts were eluted using 20%, 40%, 60%, 80%, and 100% methanol in $H_2O$ (200 ml each) at flow rate of 8 ml/min and the fractions were collected for every 200 ml. For bioassay, 5 ml were collected from each fraction, which were evaporated separately. The crude from 5 ml was dissolved in 200 µL of DMSO and 2 µL was used for bioassay by SEM imaging. The 20% methanol fraction contained active compound(s) and was used for further purification. The active semi-purified compounds were dissolved in 0.1 ml of DMSO and subjected to HPLC (Agilent1260 system) equipped with Cosmosil C18-AR-II column (Nacalai, 10 id × 250 mm). The semi-purified compounds were eluted by gradient of solvents: MeOH and 10 mM $CH_3COONH_4$ in $H_2O$. First set at 3% of methanol for 10 min and increased to 70% until 30 min. Flowrate was 3 ml/min and the column temperature was 40°C. The elution was monitored at 254 nm and the prospective peaks were collected for bioassay. Finally, 1.1 mg of a purified compound was obtained for further structure elucidation. For NMR analysis, the purified compound was dissolved in methanol-$d_4$ and $^1$H and $^{13}$C NMR spectra were collected by JMN-A500 (JEOL).

***HPLC analysis (for quantification of pyrogallol)***

Analytical HPLC was performed using HPLC (Agilent1260 system) equipped with Cosmosil PBr column (Nacalai, 2 id × 150 mm). For this, 10 µL of extracts in $H_2O$ or DMSO were subjected to HPLC and eluted by gradient mode of solvent (MeOH/0.1% formate in $H_2O$; 0–3 min: 10% MeOH, 3–14 min: 10 to 95% MeOH, 14–20 min: 95% MeOH) at a flow rate of 0.2 ml/min and the column temperature of 40°C. The elution was monitored at 265 nm and pyrogallol was eluted at 6.3 min. We used 2,3-naphthalenedialdehyde (10 µg/ml) for internal standard.

***Construction of strain expressing AcGFP1 in S. griseus***

To construct the AcGFP1 expression vector, promoter regions of *rrnD*, *hrdB* from *S. griseus* IFO13350 was amplified by polymerase chain reaction (PCR) using primers (PrrnD-F and PrrnD-R; PhrdB-F and PhrdB-R in Table S3). The gene of fluorescent protein (Acgfp1) was obtained from *Aequorea coerulescens* and amplified using pAcGFP1 plasmid (Clontech) by PCR using primers (Acgfp1-F and Acgfp1-R in Table S3). The amplified PCR fragments were tandemly connected by overlap extension PCR. The PCR product (PrrnD-PhrdB-Acgfp1) was digested by *Eco*RI and *Hin*dIII and ligated into the corresponding site of the pTYM19 (1) to construct pTYM19-AcGFP1 plasmid. Transformation of the *S. griseus* using the plasmid was performed by protoplast protocol as previously described (2).

***Image extraction of hyphae branching and processing by Fiji (ImageJ)***

The obtained CLSM images were processed using Trainable Weka Segmentation, an open-source software that combines image processing toolkit Fiji, with the data mining and machine learning toolkit Waikato Environment for Knowledge Analysis (WEKA: https://waikato.github.io/weka-wiki/) (3). We manually selected hyphae as class 1 and background as class 2. Gaussian blur, Sobel filter, Hessian, Membrane projections, and difference of gaussians were used for the training features to distinguish hyphae area and background, and FastRandomForest was used for Random Forest classifier. The classified hyphae area images were converted to binary image using Graph Cut process. The background noises were removed by Remove outlier process (Radius 2.0 pixels, Threshold 50), and skeletonization was performed using Skeletonize process. Neuroanatomy process was used to identify the order of branch (tip of hyphae) and indicated using different colors. Top ranks were identified as lengths for tip-to-branch, and others for branch-to-branch. Overlapped branches were removed manually during the process. Tip-to-branch or branch-to-branch was classified by color threshold, and the branch length was measured automatically using Analyze skeleton process. Whole microscopy images were used for the extraction process. 55 images for pyrogallol effect and 55 images for control were used for quantification analysis in FIG 5g. 58 and images for catalase effect and 38 images for control were used for quantification analysis in FIG 5h. 58 images for $H_2O_2$ effect and 31 images for control were used for quantification analysis in FIG 5i.

***Precursor (gallic acid or 2,3-DHBA) feeding experiment***

We spread 1 mL of 2,3-DHBA (3.75 mg/ml in MeOH) or gallic acid (7.5 mg/ml in MeOH) on the A3M agar medium (25 ml) and air-dried. *S. variegatus* spore stock ($10^9$ CFU), *M. septicum* cell stock ($10^{10}$ CFU), or *B. subtilis* cell stock ($10^7$ CFU) were spread on the agar medium, and cells were grown by incubation at 30°C for 2 days. After the incubation, the agar medium was extracted using HClaq (pH 2.2) and filtered (0.2 μm) for further PDA-HPLC analyses.

***Bioconversion of 2,3-DHBA to pyrogallol***

For the combination assay to elucidate the pyrogallol producer, *S. variegatus* and *M. septicum* was precultured using ISP2 medium (10 ml in test tube) at 30°C for 2 days. *B. subtilis* was precultured by Miller Hinton medium (10 ml in test tube) at 30°C for 2 days. Preculture of *S. variegatus* (50 μL) and *M. septicum* (50 μL), *B. subtillis* (15 μL) was used to inoculate the A3M agar medium (15 ml) and further incubated at 30°C for 2 days. After the incubation, agar medium was crushed by glass syringe and 15 ml of 6.3 mM HCl (pH .2.2) was added to extract the metabolites. After centrifugation (12,000 rpm, 15 min), 100 μl of 2,3-naphthalenedialdehyde (0.1 mg/ml) was added to 900 μL of extracts as internal standard for HPLC quantification. A total of 10 μl was subjected for further HPLC analyses. For the mixed culture of *M. septicum* with knockout mutants of *dhb* genes (for bacillibactin biosynthesis) (Δ*dhbA*, Δ*dhbB*, Δ*dhbC*, Δ*dhbE*, and Δ*dhbF*), 168 wild-type and the mutants of *B. subtilis* were precultured by Miller Hinton medium (10 ml in test tube) until $OD_{660}$ reach 0.4 and then 15 μL was used to inoculate the A3M agar medium (15 ml) and further incubated at 30°C for 5 days. Pyrogallol was extracted as same procedure as described above.

***Heterologous expression of nahG from M. septicum***

Chromosomal DNA of *M. septicum* was obtained using CTAB method (4). Short read sequences obtained using Hi-seq (Illumina) were associated with contigs, and CDS were deduced using

RAST pipeline (https://rast.nmpdr.org/) (5) to search for NahG homologs. The DNA sequence of *nahG*$^{Ms}$ containing fragment was registered to the database with accession number LC666909. Thereafter, the DNA fragment that codes *nahG* gene was amplified by PCR using primers: MsnahG-F and MsnahG-R (Table S3). The DNA fragment was digested by *Nde*I and *Hin*dIII and ligated into the corresponding site of pET26b to generate pET26-nahG$^{Ms}$ to express NahG$^{Ms}$ with no additional tags. pET26-nahG$^{Ms}$ was used to transform *Escherichia coli* BL21(DE3) and cultured using LB with 50 µg/ml kanamycin at 37°C. Preculture was performed by 4 ml LB at 37°C, 200 rpm for 1 day, and this preculture was used to inoculate 100 ml LB with 50 µg/ml kanamycin and 1 mM IPTG in 500 ml flask at 37°C and 200 rpm for 1 day. Thereafter, it was centrifuged (3,000×g, 4°C) to obtain the cells. The cell pellets were washed twice using 20 ml of 0.1 M sodium phosphate buffer (pH6.8). Thereafter, 40 ml of sodium phosphate buffer was added to suspend the cells and 995 µl of cell suspension was transferred into 1.5 ml tube to prepare the resting cell. For the bioconversion experiment, 5 µl of 40 mM 2,3-DHBA in methanol was added to the resting cells, and 5 µl of 40 mM salicylic acid in ethanol was added to the resting cells for positive control. *E. coli* BL21(DE3) with empty vector was used as negative control. The reaction samples were incubated at room temperature for 9 hrs at 1,200 rpm. The reaction mixture was centrifuged (20,400×g, 10 min, 4°C) and the supernatant (5 µL) was subjected to HPLC analysis. HPLC condition for pyrogallol detection is described above.

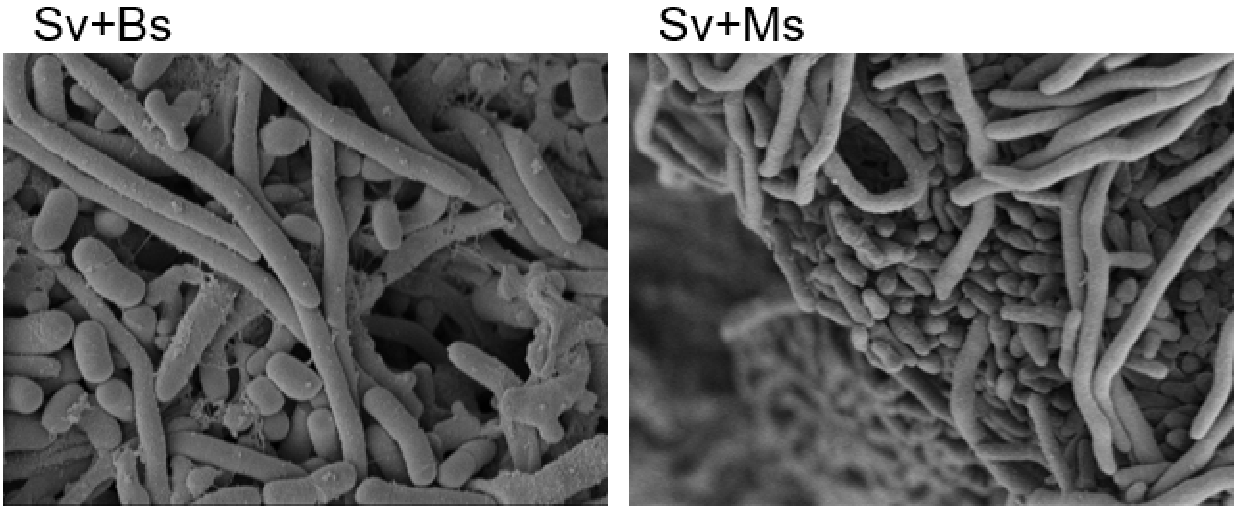


**FIG S1** Scanning electron microscope (SEM) images of cell morphology in competitive dual culture. Sv: *Streptomyces variegatus* HEK138A, Ms: *Mycolicibacterium septicum* HEK138M, Bs: *Bacillus subtilis* 168.

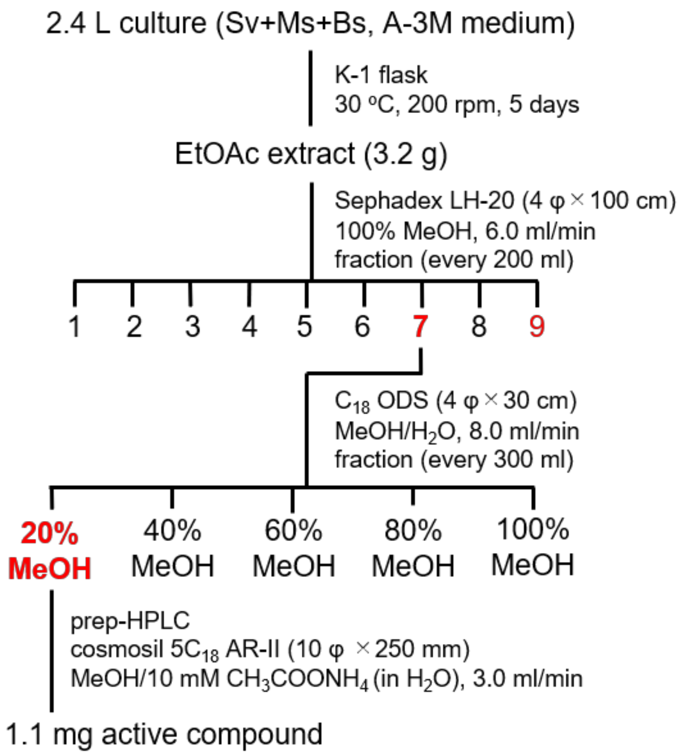


**FIG S2** Purification scheme of active compound. Sv: *S. variegatus* HEK138A, Ms: *M. septicum* HEK138M, Bs: *B. subtilis* 168.

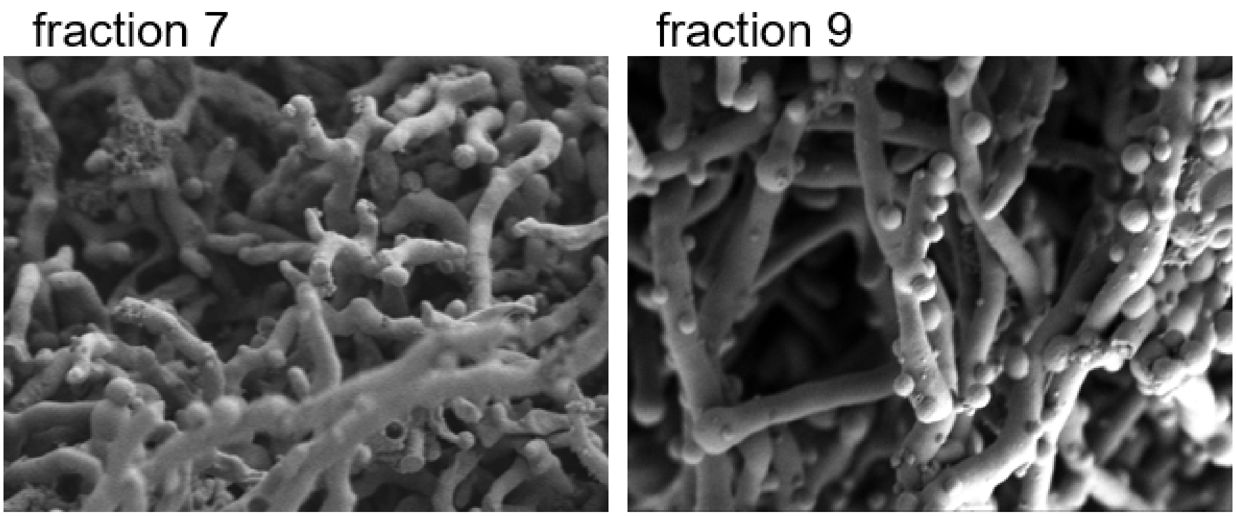


**FIG S3** SEM images of *S. variegatus* HEK138A mycelia treated by active fractions. SEM images of *S. variegatus* mycelia treated by active fractions No. 7 and No. 9 from Sephadex LH-20 column.

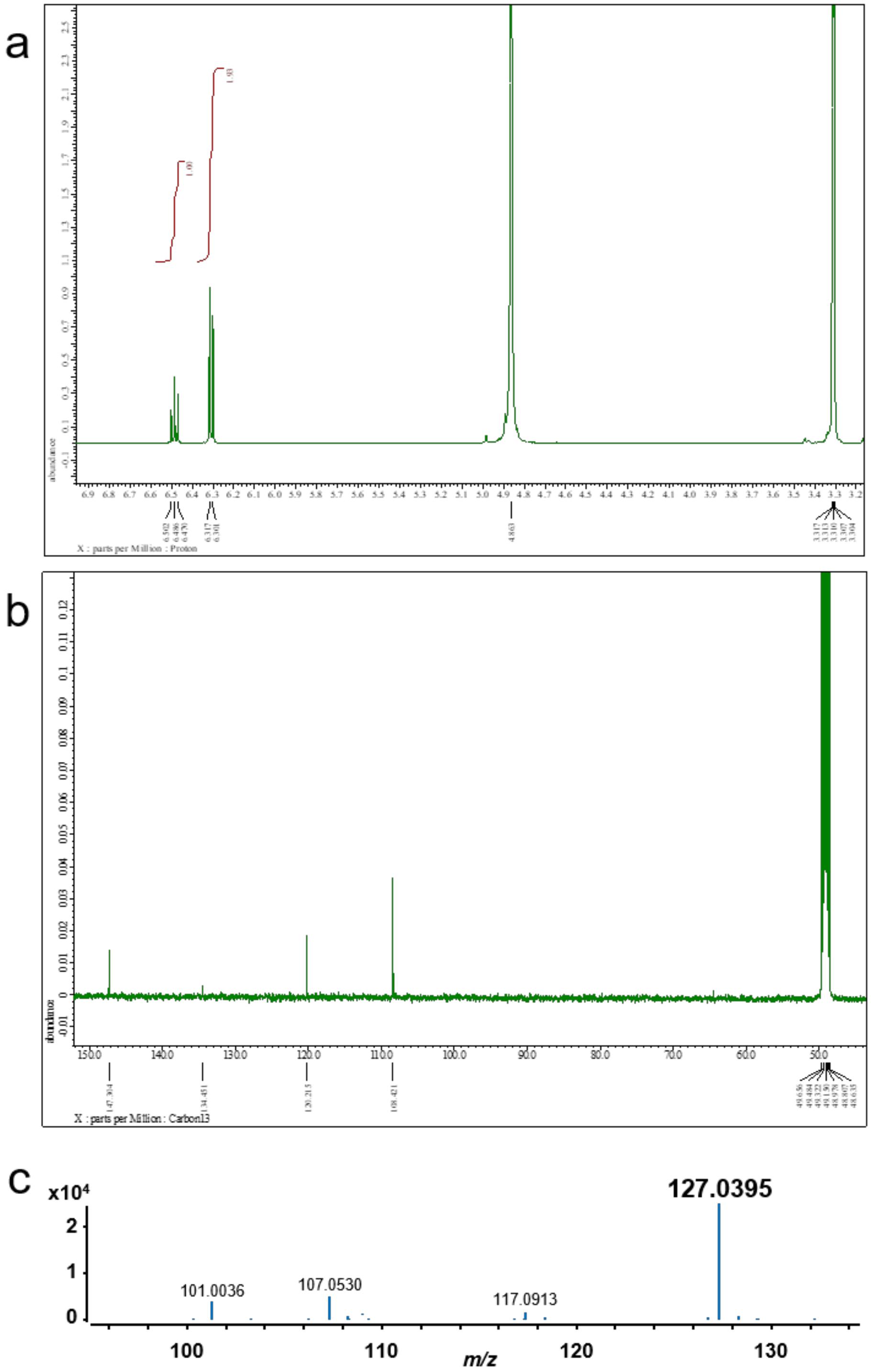


**FIG S4** NMR and MS analysis of the isolated active compound. (a) $^{1}$H NMR (500 MHz) of isolated pyrogallol in methanol-$d_4$. (b) $^{13}$C NMR (125 MHz) of isolated pyrogallol in methanol-$d_4$. The methanol-$d_4$ signal ($^{1}$H: 3.31 ppm, $^{13}$C: 49.15 ppm) was used as a reference. (c) HR-ESI-QTOF-MS of isolated pyrogallol.

**TABLE S1** NMR assignments.

| position | δC | δH |
|---|---|---|
| 1 | 134.5 | |
| 2 | 147.3 | |
| 3 | 108.4 | 6.31 (2H, d, 8.0) |
| 4 | 120.2 | 6.49 (1H, t, 8.2) |

methanol-$d_4$ signal ($^{1}$H: 3.31 ppm, $^{13}$C: 49.15 ppm) was used as a reference.

**TABLE S2** MIC of pyrogallol against tested microorganism.

| Strain | Pyrogallol (μg/ml) |
|---|---|
| *Streptomyces variegatus* HEK138A | 32 |
| *Mycolicibacterium septicum* HEK138M | 32 |
| *Bacillus subtilis* 168 | 256 |
| *S. coelicolor* A3(2) | 32 |
| *S. lividans* TK23 | 32 |
| *S. avermitilis* MA-4680 | 32 |
| *S. griseus* IFO13350 | 32 |
| *Staphylococcus aureus* 209PJC-1 | 32 |
| *Micrococcus luteus* ATCC9341 | >512 |
| *B. subtilis* ATCC6633 | 128 |
| *Escherichia coli* NIHJ-2 | 128 |
| *Saccharomyces cerevisiae* | 64 |
| *Candida albicans* | >512 |

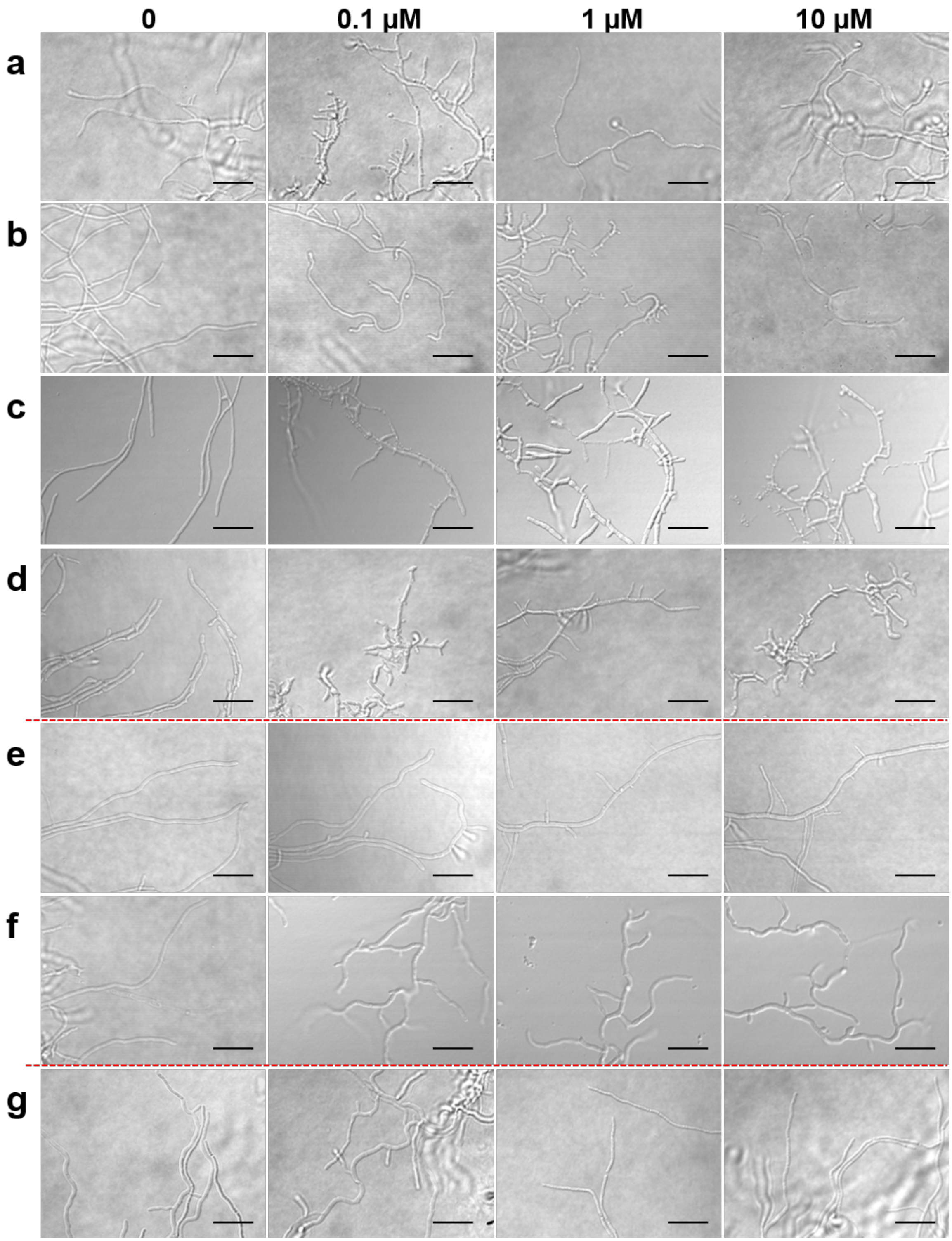


**FIG S5** DICM images of *Streptomyces* spp. mycelium treated with 0.1-10 μM of pyrogallol. (a) *S. variegatus* HEK138A; (b) *S. griseus* IFO13350; (c) *S. albus* J1046; (d) *S. lividans* TK23; (e) *S. venezuelae* NBRC13096; (f) *S. avermitilis* MA-4680; (g) *S. coelicolor* A3(2). (scale bar = 5 μm)

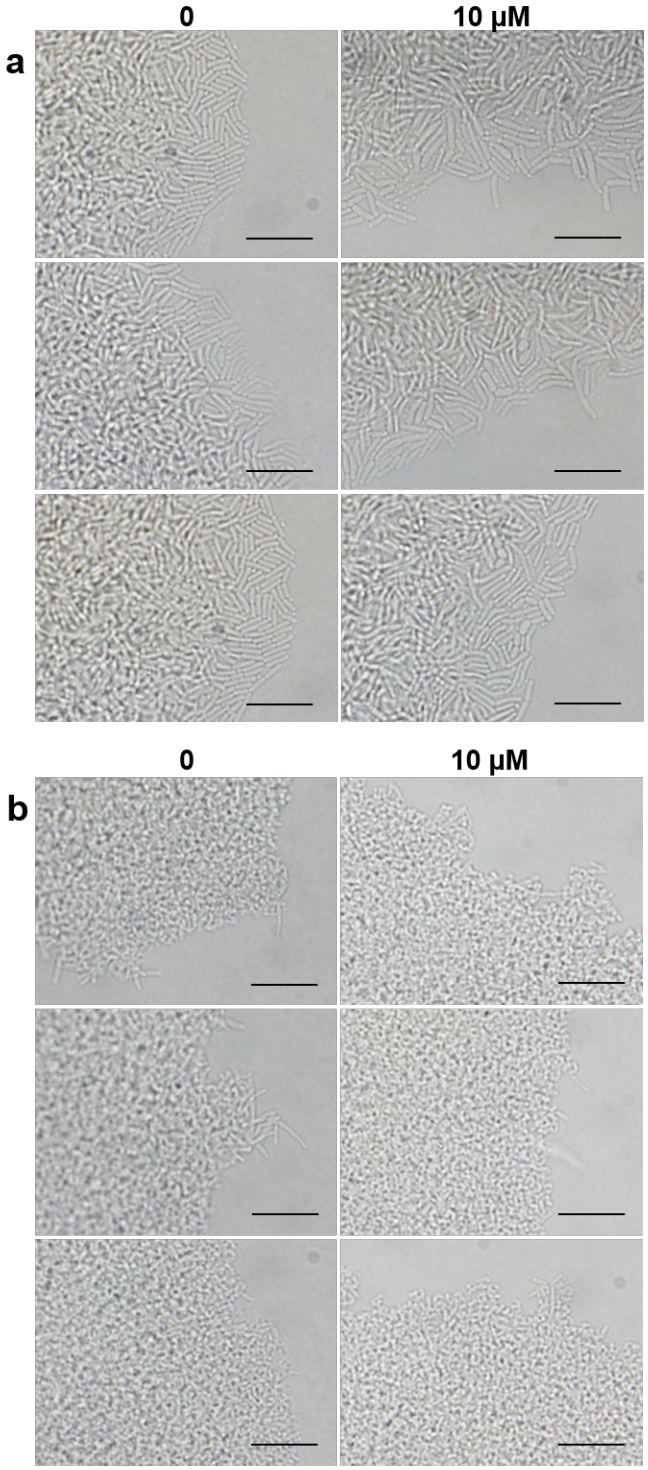


**FIG S6** DICM images of *B. subtilis* cells (a) and *M. septicum* HEK138M (b) treated with pyrogallol (selected three images for each). Left: no addition of pyrogallol. Right: Addition of 10 μM of pyrogallol. (scale bar = 5 μm)

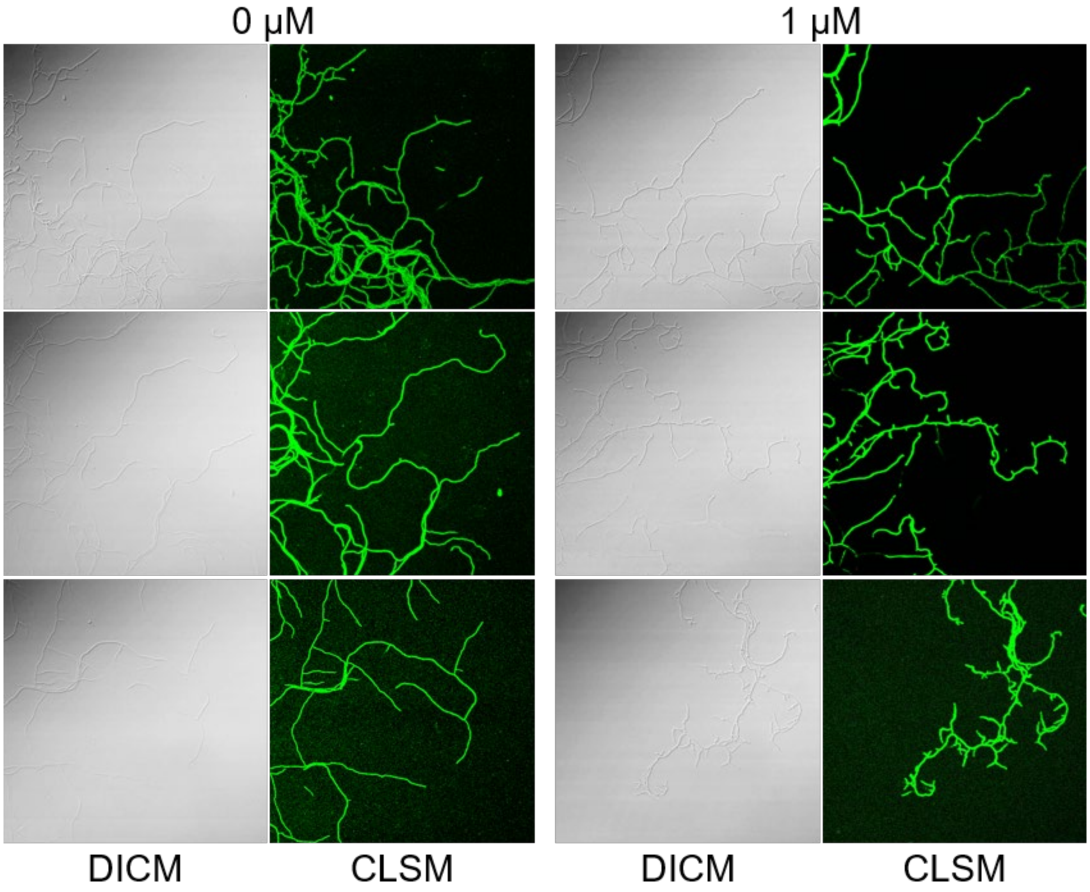


**FIG S7** DICM and CLSM images of *S. griseus* expressing AcGFP1 treated with pyrogallol (selected three images for each). Left: no addition of pyrogallol. Right: Addition of 1 μM of pyrogallol.

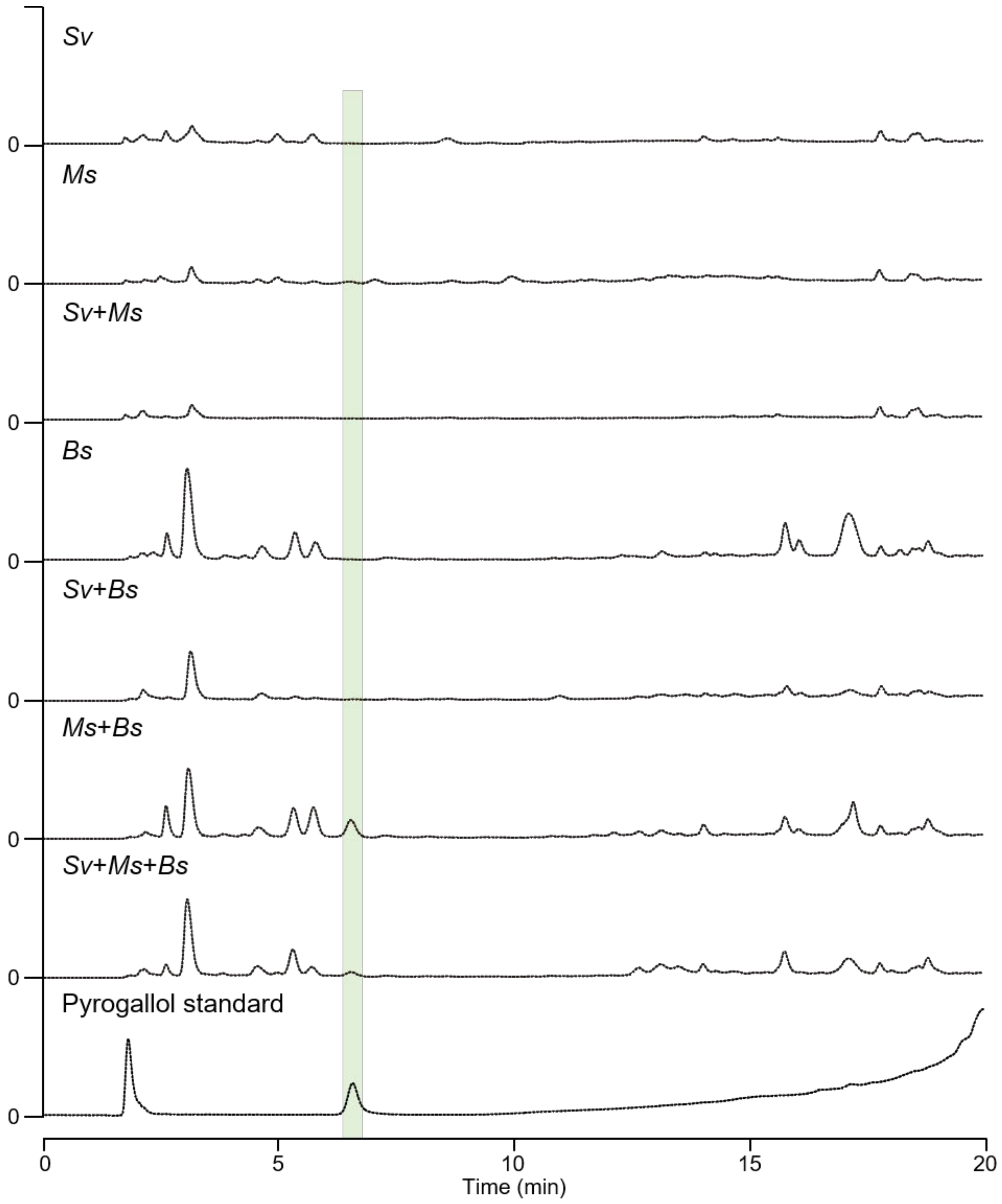


**FIG S8** PDA-HPLC traces of metabolites in combination of all mixed-culture. Sv: *S. variegatus* HEK138A, Ms: *M. septicum* HEK138M, Bs: *B. subtilis* 168. Pale green background indicates the peak of pyrogallol.

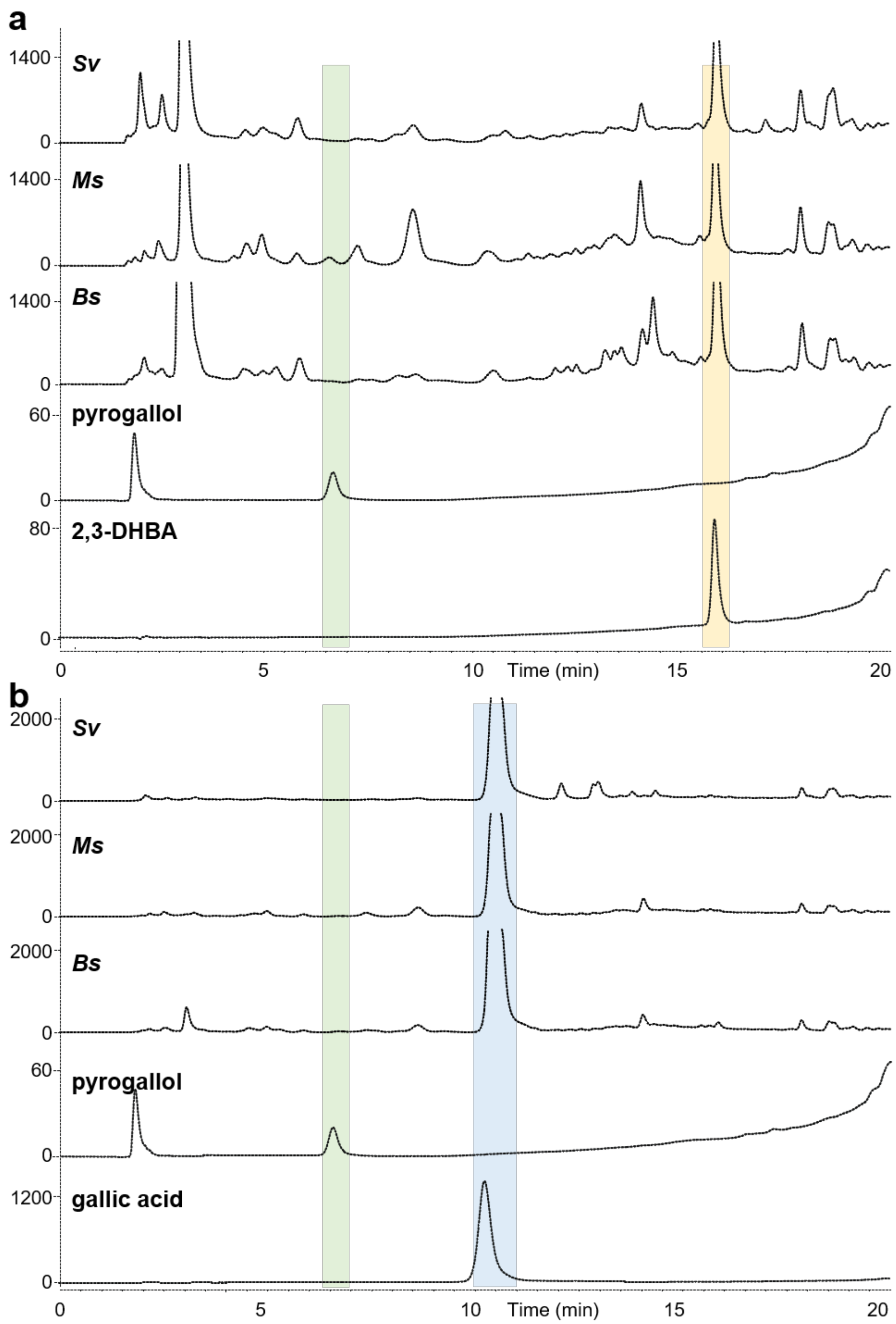


**FIG S9** PDA-HPLC traces of 2,3-DHBA (a) or gallic acid (b) feeding experiment. Sv: *S. variegatus* HEK138A, Ms: *M. septicum* HEK138M, Bs: *B. subtilis* 168. Pale green, pale yellow, and pale blue backgrounds indicate the peaks of pyrogallol, 2,3-DHBA, and gallic acid, respectively.

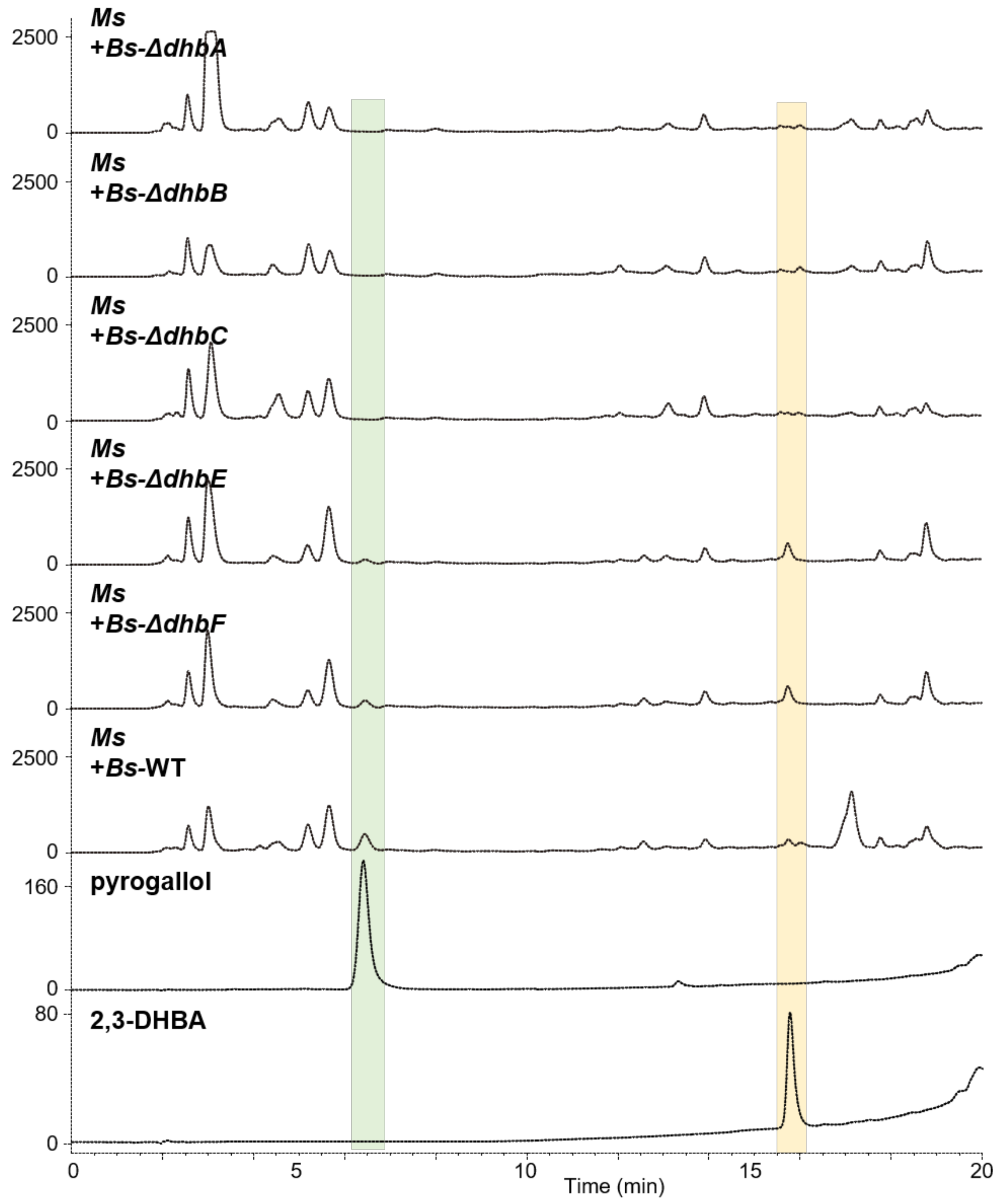


**FIG S 10** PDA-HPLC traces of co-culture between Ms and *dhb* gene knockout mutants. Ms: *M. septicum* HEK138M, Bs: *B. subtilis* 168. Pale green and pale yellow backgrounds indicate the peaks of pyrogallol and 2,3-DHBA, respectively.

```
NahG            MQNSTSALNVSIIGGGIAGVALALDLCRHAHLNVQLFEAAPAFGEVGAGVSFGANAVRAI
NahG-Ms         ----MTGLRVAVVGAGIGGLTAAIALRANG-IDATVYEQAHELKALGAGVAIATNGSRIL
                     :.*.*:::*.**.*:: *: *  :. ::. ::* *  :  :****::.:*. * :

NahG            AGLGIAEPYGKIADSNPAPWQDIWFEWRNGRDAKYLGCSVAEGVGQSSVHRADFLDALAS
NahG-Ms         NKLGVGDAVAAIAGPVTHYQFRTWQAEPIAGEPSTLG--FGDPARTWCLHRGEFQKVLAD
                  **:.:. . **.. .      *     . :.. **  ..: .   .:**.:* ..**.

NahG            QLPDGIAQFGKRAQRVEQDGEQVRVTFTDGSEHRCDLLIGADGIKSSIRDHVLQGLNQPL
NahG-Ms         ALPVDALQLGRSCVGATEYGDGVRVEFSDGTTVDADLLVGADGIHSRLQGKVTR-----P
                 ** .  *:*: .  . : *: *** *:**:   .***:*****:* ::.:* :

NahG            ASPRFSGTCAYRGLIDSQQLREAYRARGVDEHLIDVPQMYLGLDGHILTFPVKQGRLINV
NahG-Ms         AAPVSEGIMAYRGLIPADRLRGVVDMN--------ASSMWLGPRQSFLAYPVSAGELINI
                *:*  .*  ****** :::** .   .        ...*:**    :*::**. *.***:

NahG            VAFISDRSQPNPVWPSDTPWVRNATQAEMLAAFEGWDDAAQVLLECIPTPSLWALHDLAE
NahG-Ms         VAFVPTN------LTVTESWTAPGDVAELAAAYRGWDPRVSSIIDAMDSTFRWGIYDREP
                ***:. .       .    .*.  .  **: **:.***  .. :::.: :.  *.::*

NahG            LPGYVHGRVVLIGDAAHAMLPHQGAGAGQGLEDAWLLARLLEDPKVLDKRPQAVLDAYDA
NahG-Ms         LDRWSTDRITLLGDSAHAVTPHLGQGANQAIEDAMTLAVVLRDAQPGEIG--TRLRHYEN
                *  :  .*:.*:**:***: ** * **.*.:***  ** :*.*.:  :    : *  *:

NahG            VRRPRACRVQRTSFEAGELYEFRDPAVLADEERLGKVLAERFDWLWNHDMQEDLLQAREL
NahG-Ms         LRIDRTGHVRRQARAAGHIYRSTELTPHAQAEQLQAILDS---------VAINTYDAERV
                :*  *: :*:* :  **.:*.  : :  *: *:*  :* .         :  :  :*..:

NahG            LGLRAQAA
NahG-Ms         AENAGQAA
                    .***
```

**FIG S11** ClustalW alignment of NahG from *Pseudomonas putida* and *M. septicum* HEK138M. ClustalW pairwise alignment was performed in website (https://www.genome.jp/tools-bin/clustalw). Asterisk (*) indicates identical amino acids. Colon and period (: and .) indicate similar amino acids.

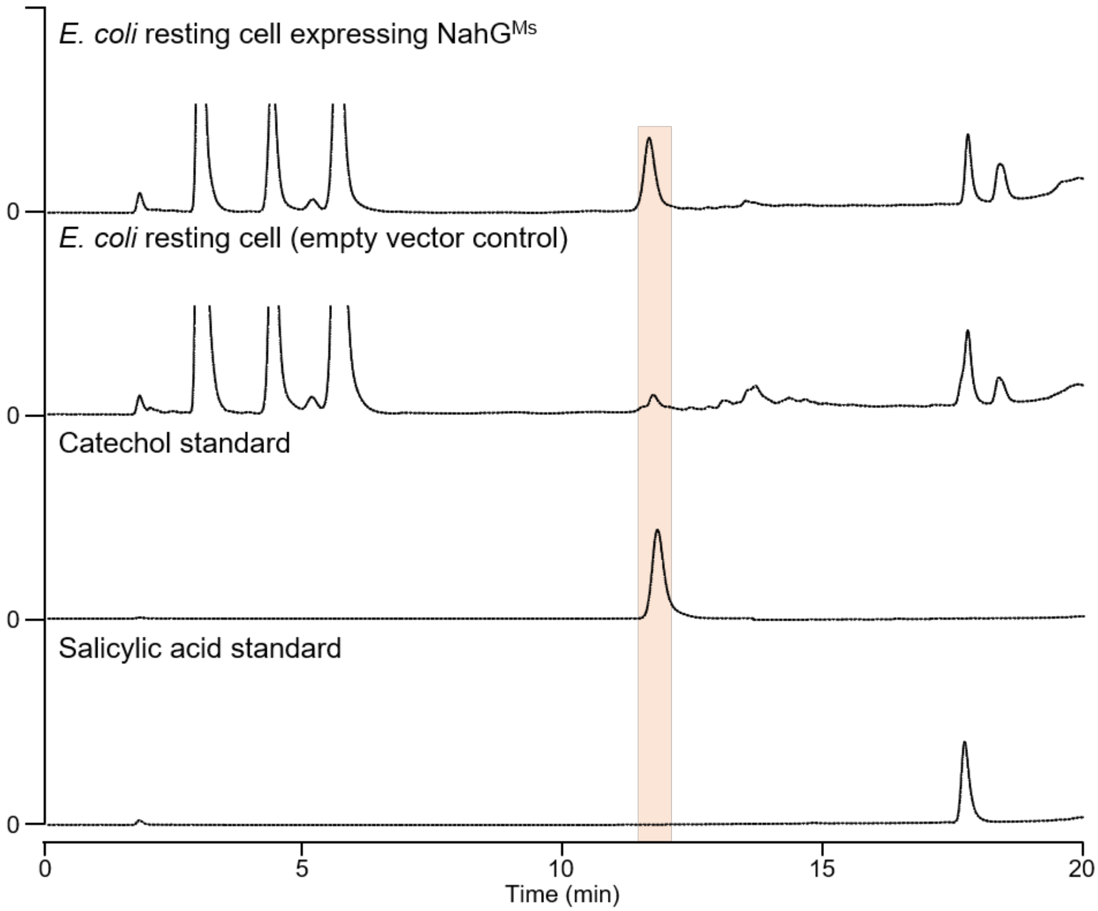


**FIG S12** Bioconversion of salicylic acid to catechol by *E. coli* resting cell expressing NahG$^{Ms}$. PDA-HPLC traces of salicylic acid to catechol. Catechol is indicated in pale red background.

**TABLE S3** Oligo-DNA primers used for PCR in this study.

| Primer name | Primer sequence (5' to 3') |
|---|---|
| PrrnD-F | CGC AAG CTT GCT TTC TCG CGT ATG TC |
| PrrnD-R | TTC CGC ACG CTC ACC ATG TTT ACC CGT AAT CGG T |
| PhrdB-F | GGT GAG CGT CGC GAA GGA A |
| PhrdB-R | CGC CCT TGC TCA CCA TGA ACA ACC TCT CGG AAC G |
| Acgfp1-F | ATG GTG AGC AAG GGC GCC GAG |
| Acgfp1-R | GCG AAT TCT CAC TTG TAC AGC TCA TCC A |
| MsnahG-F | CCC AGA TAT ACA TAT GAC CGG ACT TCG AG |
| MsnahG-R | TTT CGC AAG CTT TCA GGC G |
